# *Leishmania amazonensis* sabotages host cell SUMOylation for intracellular survival

**DOI:** 10.1101/2021.11.03.467107

**Authors:** Kendi Okuda, Miriam Maria Silva Costa Franco, Ari Yasunaga, Ricardo Gazzinelli, Michel Rabinovitch, Sara Cherry, Neal Silverman

## Abstract

*Leishmania* parasites use elaborate virulence mechanisms to invade and thrive in macrophages. These virulence mechanisms inhibit host cell defense responses and generate a specialized replicative niche, the parasitophorous vacuole. In this work, we performed a genome-wide RNAi screen in *Drosophila* macrophage-like cells to identify host factors necessary for *Leishmania amazonensis* infection. This screen identified 52 conserved genes required specifically for parasite entry, including several components of the SUMOylation machinery. Further studies in mammalian macrophages found that *L. amazonensis* infection inhibited SUMOylation within infected macrophages and this inhibition enhanced parasitophorous vacuole growth and parasite proliferation through modulation of multiple genes especially *ATP6V0D2*, which in turn effects *CD36* expression and cholesterol levels. Together, these data suggest that parasites actively sabotage host SUMOylation and alter host transcription to improve their intracellular niche and enhance their replication.

## Introduction

*Leishmania* parasites use multiple strategies to thrive in mammalian phagocytes. One striking example is the use of virulence factors to evade or down regulate the host immune response. For example, amastigote forms of the parasite invade macrophages, and express surface molecules that engage phagocytic receptors that promote ‘silent’ or anti-inflammatory cell invasion, typically observed with the engulfment of apoptotic cells [1–5]. Inside macrophages, amastigotes live within specialized replicative organelles known as parasitophorous vacuoles (PVs), which are modified mature phagosomes. Amastigotes create a hospitable niche in the PV through secretion of virulence factors that dampen host cell defenses. These secreted factors work through many mechanisms including altering host protein expression and modulating post-translational protein modifications [6, 7, reviewed in 8, 9-13]. Our understanding of the virulence mechanisms used by these parasites is incomplete while a deeper understanding is essential to develop new strategies to combat leishmaniasis, which threatens 0.9 million people annually [14].

Mammalian models of *Leishmania* infection have been essential to build our current understanding of leishmaniasis. However, other models of infection, such as *Drosophila melanogaster*, are attractive alternatives because they share many key signaling pathways with mammals and have powerful genetic and genomic techniques readily accessible. Here, we exploited the *Drosophila* SL2 macrophage-like cell line in a genome-wide RNAi screen to identify host factors required for *L. amazonensis* infection. With this screen, we identified and validated 52 conserved host factors specifically involved in *L. amazonensis* entry but not *Escherichia coli* nor *Staphylococcus aureus* phagocytosis. These hits were particularly enriched in SUMOylation factors.

SUMOylation is a ubiquitous post-translational modification in which a Small Ubiquitin-like MOdifier protein (Sumo) is conjugated to lysine (Lys) residues on target proteins, through the sequential action of E1 activating (Sae1/Uba2 dimer), E2 conjugating (Ubc9) and E3 ligases (several of them) enzymes [15]. SUMOylation regulates gene expression, protein function, localization or activity [16], and 6,747 different SUMOylated proteins were cataloged in HeLa and U2OS cell lines [17].

Interestingly, many pathogens, including virus, bacteria and protozoans, target SUMOylation to exploit host cellular functions or to inhibit cell defense responses [18-23]. Here, we observed that intracellular *L. amazonensis* amastigotes efficiently inhibited overall macrophage SUMOylation, reducing this post-translational modification to nearly undetectable levels. Moreover, SUMO1 or SUMO2 depleted cells presented enhanced parasite growth with larger PVs, higher expression of *ATP6V0D2*, and consequently elevated *CD36* and cholesterol levels. Therefore, our data suggest that inhibition of SUMOylation is a novel virulence mechanism of *L. amazonensis* parasites, used for successful infection of macrophages.

## Results

### The *Drosophila* cell model of *L. amazonensis* infection

*Drosophila* macrophages, also known as plasmatocytes, play key roles in immune defense and development [24]. Previously, we demonstrated that these macrophages use CD36-like scavenger receptors to phagocytose and control proliferation of *L. amazonensis* in the hemolymph of adult flies [25]. In this work, we determined if the *Drosophila* macrophage-like SL2 cell line would phagocytose parasites, in particular the amastigote form that is responsible for repeated cycles of macrophage infections in mammalian hosts. SL2 cells were infected with antibody free *L. amazonensis* amastigotes which were promptly phagocytosed and observed within SL2 cells (Fig 1A). By 48 h post infection, parasites were inside small acidic (Lysotracker positive) vacuoles (Fig 1B), and by 96 h post infection intracellular parasites were mostly cleared from these cells (not shown).

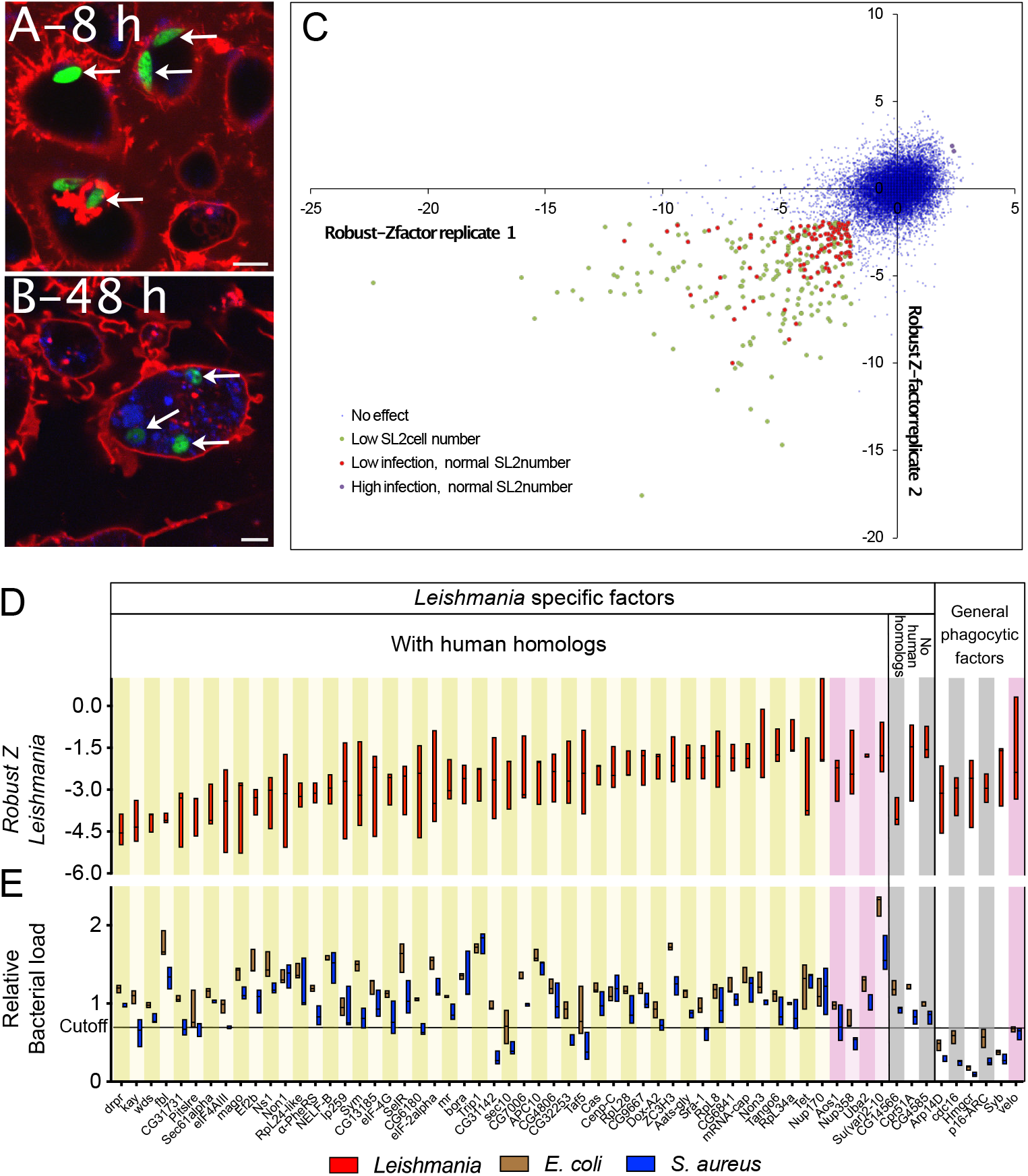

As PV biogenesis and parasite proliferation were not observed in the SL2 cell model, this allowed us to specifically focus on the phagocytosis and entry of amastigotes, a step that is essential for leishmaniasis development. To quantify intracellular parasites, SL2 cells were infected with GFP-expressing amastigotes, fixed and extracellular parasites were immunostained with anti-*Leishmania* serum. The ratio of intracellular parasites (immunostaining negative) and SL2 cell number (Hoechst 33342 stain of nucleus) was scored microscopically using image analysis software (Figure S1A, B). The phagocytosis of parasites increased over the 4 h time course of this assay, with ~70% of the internalization completed by 90 min (Fig S1C). The effect of several multiplicities of infection was analyzed at 90 min post infection (pi). While the total number of internalized parasites increased with higher MOI (Fig S1D), parasite aggregation also increased, making reliable quantification difficult (Fig S1E). Therefore, we selected a MOI of 0.5 for 90 min for screening. We tested the robustness of the assay by infecting and scoring 24 replicate wells, either with no RNAi, or transfected with RNAi targeting a control (non-silencing) gene, or targeting genes known to be required for phagocytosis (βCOP, Cdc42) [25–29] (Fig S2A-B). These phagocytosis genes clearly and significantly reduced amastigote entry.

We then used this assay to identify factors required for the internalization of *L. amazonensis* into SL2 cells by performing a genome-wide RNAi screen with a library of ~13,000 dsRNAs covering nearly the entire *Drosophila* genome (Ambion). The number of intracellular parasites per cell for each RNAi treatment was quantified and the Robust Z-score was calculated (the distance of the infection in each well from the median of the entire plate) [30].

The entire library was screened in two completely independent replicates and the Z-scores of infection from the duplicate screens are presented in the Figure 1C, where X- and Y-axes display the Z-scores of each replicate, and each dot represents a distinct RNAi treatment (the complete dataset of preliminary hits is also presented in Table S1). We used a cutoff of −≤−2.0 or ≥2 in both replicates (p<0.001) to identify the genes required for infection and genes that restrict infection (referred to as ‘down ‘and ‘up’ hits hereafter). Further, those candidates that reduced SL2 cell viability and/or proliferation (green dots in Figure 1C) were removed from further analysis. This analysis revealed 120 candidate genes (118 downs and 2 up hits, red and purple dots in Figure 1C). Independent dsRNAs targeting these candidates, free of predicted off-targets [31], were synthesized, and used in validation assays performed in triplicate. This secondary screen validated 61 down hits but neither of the initial candidate ups (Figure 1D; Table S1).

Amongst these 61 down hits, we next sought to determine which generally impact phagocytosis and which are specifically required for uptake of *Leishmania* amastigotes.

To this end, we performed phagocytosis assays with FITC-labeled *E. coli* and *S. aureus* [32]. We quantified uptake of FITC labeled *E. coli* and *S. aureus* using flow cytometry [29]. Six candidates reduced the internalization of both *E. coli* and *S. aureus* by ≥30% and were thus classified as general phagocytic factors, while 55 were more specific to *Leishmania* (Fig. 1E, and Table S1). Three of the *Leishmania* specific hits have no clear human homologs, leaving 52 down hits for further investigations.

### *Leishmania* uptake factors are highly enriched in protein SUMOylation

In order to identify host cell biological processes affected by the 52 conserved hits from our genome-wide RNA screen, these candidates were submitted to an overrepresentation test using Panther, with the *Drosophila* GO network [33]. “Protein SUMOylation” was 99-fold enriched with a P-value of 8×10^-6^ (Table 1). Three of the eight genes that make up the protein SUMOylation group were hits - both SUMO E1 subunit genes *Aos1* and *Uba2*, as well as the SUMO E3 ligase, *Su(var)2-10* (aka PIAS). Additionally, the single *Drosophila* SUMO ortholog *smt3* reduced infection, but also affected cell proliferation, while the SUMO protease *velo* (SENP6/7 homolog) affected both *Leishmania* entry and bacterial phagocytosis (Figure 1D-E marked with pink stripes, and Table S1).

| *Drosophila melanogaster* |  |  |  |  |  |
| --- | --- | --- | --- | --- | --- |
| GO biological process complete | Total genes # | From input # | Fold Enrichment | raw P value | Genes |
| protein sumoylation | 8 | 3 | 99.28 | 8.12E-06 | Su(var)2-10, Aos1, Uba2 |
| ribosomal large subunit assembly | 20 | 4 | 52.95 | 1.79E-06 | Non3, CG13185, RPL24, CG11583 |
| mitotic spindle assembly | 25 | 3 | 31.77 | 1.54E-04 | Non1, Non3, Ns1 |
| positive regulation of Toll signaling pathway | 31 | 3 | 25.62 | 2.77E-04 | Pitslre, Aos1, Uba2 |
| positive regulation of multi-organism process | 146 | 6 | 10.88 | 2.15E-05 | Pitslre, Su(var)2-10, Sec61alpha, Aos1, kay, Uba2 |
| translation | 307 | 10 | 8.62 | 2.36E-07 | GlyRS, RpL28, EF2, eIF2alpha, RpL8, eIF3i, RpL24, RpL34, eIF4G1, alpha-PheRs |
| RNA metabolic process | 828 | 12 | 3.84 | 5.07E-05 | Tet, GlyRS, Taf5, mago, Non3, CG7483, mRNA-cap, CG13185, CG6841, CG9667, Sym, alpha-PheRS |

### *L. amazonensis* drastically reduces macrophage SUMOylation

We next set out to define a role for protein SUMOylation in *Leishmania* infection of mammalian macrophages. We infected mouse bone marrow derived macrophages (BMDMs) with *L. amazonensis* amastigotes and harvested whole cell lysates at various time points after infection; total protein SUMOylation was analyzed by immunoblotting for SUMO1 or SUMO2/3 (Figure 2A, B, left). Uninfected macrophages presented numerous SUMO-conjugated bands ranging from ~60 to >250 kDa. This overall SUMOylation signal visibly decreased 2 h after *L. amazonensis* amastigote infection and was nearly undetectable by 48 h post infection, with either SUMO1 or SUMO2/3 probes. The infection did not change the quantity of free SUMO2/3 bands (Figure 2A, 15 kDa band), free SUMO1 was not detectable. This loss of SUMOylation was not due to total protein degradation or loss, as normal levels of GAPDH and total protein were observed by immunoblotting or Coomassie Blue staining, respectively (Figure 2 at bottom of each blot and S3).

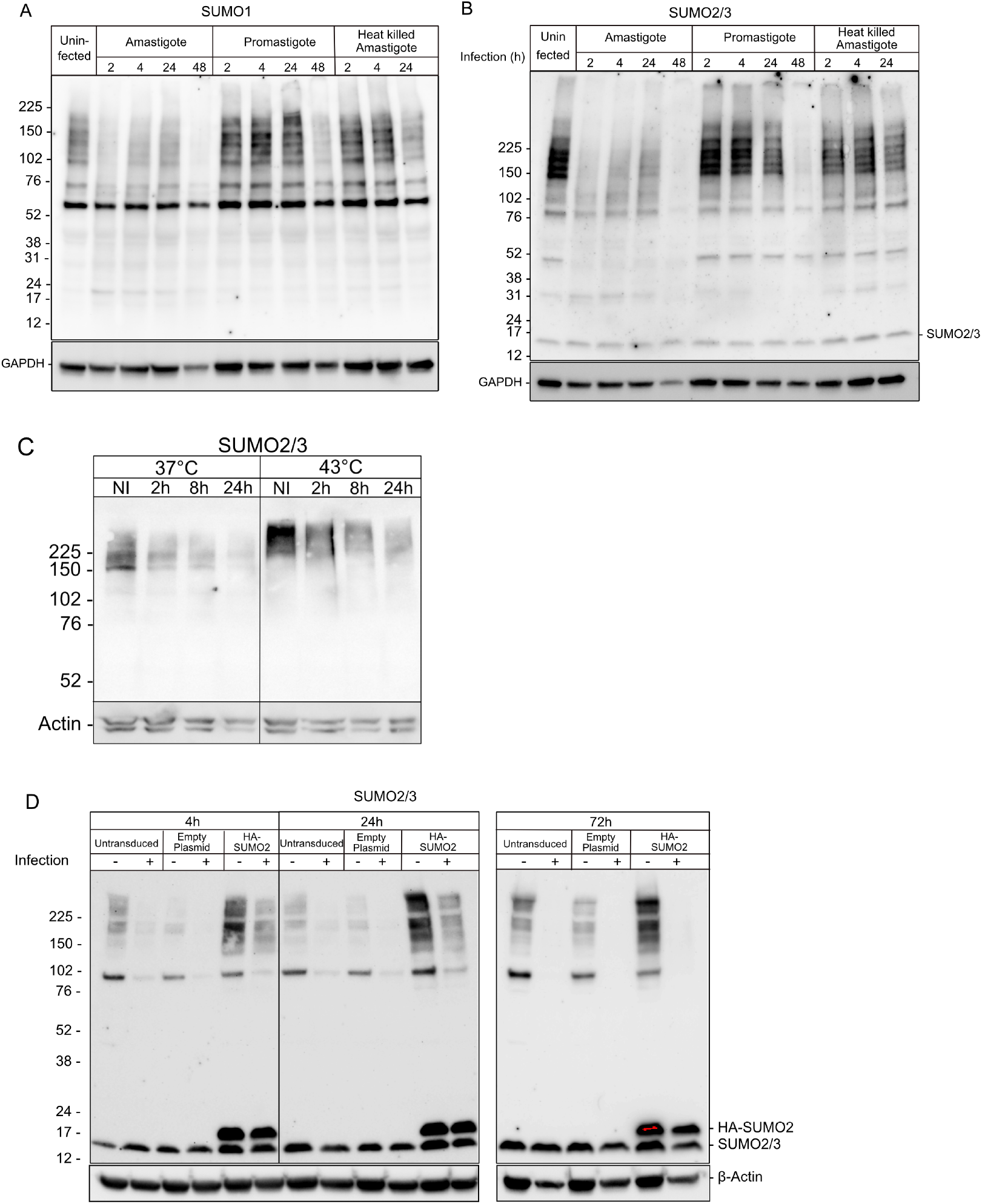

The sandfly bite transmits the promastigote form of the parasite to human skin, where they infect phagocytes by receptor mediated phagocytosis [34]. Once internalized, promastigotes differentiate into amastigotes, a differentiation process that typical requires 48 h - 72 h [35]. Interestingly, BMDM infection with promastigotes caused a delayed reduction of SUMO1 and SUMO2/3 conjugation, which was observed only after 48 h of infection (Figure 2A, B, middle). This delayed inhibition of SUMOylation with promastigote infection correlates with the time required for differentiation of promastigote into amastigotes [35].

Phagocytosis of heat-killed amastigotes slightly reduced SUMO1ylation at 24 h but no differences were observed for SUMO2/3 (Figure 2A-B, right). All together, these data argue that live intracellular amastigotes, but not promastigotes, are potent inhibitors of overall macrophage protein SUMOylation.

Next, we investigated if the intracellular parasites were able to inhibit protein SUMOylation induced by heat shock, a classic inducer of elevated SUMOylation [36]. BMDMs were infected with amastigotes for various times, and then were heat shocked at 43°C for 30 min. Similar to observations in the human lymphoblast cell line K-562 [37], heat shock induced the SUMOylation of high molecular weight proteins in uninfected BMDMs (Figure 2C, NI). However, this SUMOylation was strongly inhibited in cells infected with amastigotes for 2 h, 8 h or 24 h (Figure 2C), indicating that *Leishmania* parasites actively suppress SUMOylation induced by external stimuli.

To determine if similar regulation of SUMOylation occurs in human systems, THP-1 macrophages [38] were similarly analyzed following amastigote infection. The infection of these human macrophages with amastigotes drastically reduced overall levels of protein SUMO2/3-ylation with little or no change in free SUMO2/3 (Figure 2D). The overall protein SUMO2/3ylation was at undetectable levels at 72 h post infection. In addition, a THP-1 cell line was generated that constitutively over-expresses HA-SUMO2, which exhibited increased levels of free SUMO2 as well as SUMOylated proteins in uninfected cells (Figure 2D). While the loss of SUMOylation was delayed in this case, parasite infection still reduced it to undetectable levels by 72 hours post infection.

### SUMOylation inhibition favors *Leishmania* infection

Several pathogens sabotage the SUMOylation machinery to better survive in the vertebrate host [39–41]. We hypothesized that protein SUMOylation attenuates amastigote proliferation, and the inhibition of SUMOylation by *Leishmania* amastigotes may enhance their survival or replication. To evaluate this hypothesis, THP-1 cells were transduced with lentivirus stably expressing shRNAs specifically targeting SUMO1 or SUMO2. The efficiency of knockdowns was approximately 50%, determined by qRT-PCR (Figure 3A). The initial number of parasites per cell was not significantly affected, indicating amastigote entry was unaffected (Figure 3B, 6 h post infection). However, by 72 h the number of parasites per infected cell was significantly higher in SUMO2-knock down cells compared to the control cells (Fig 3B). SUMO1 knockdown trended in the same direction but did not reach statistical significance. These data demonstrate that *L. amazonensis* replicates more effectively in macrophages with reduced SUMOylation capacity. These findings are consistent with our hypothesis that SUMO-dependent process interfere with amastigote growth and argue that amastigotes restrain macrophage SUMOylation to enhance their intracellular niche and increase parasite growth.

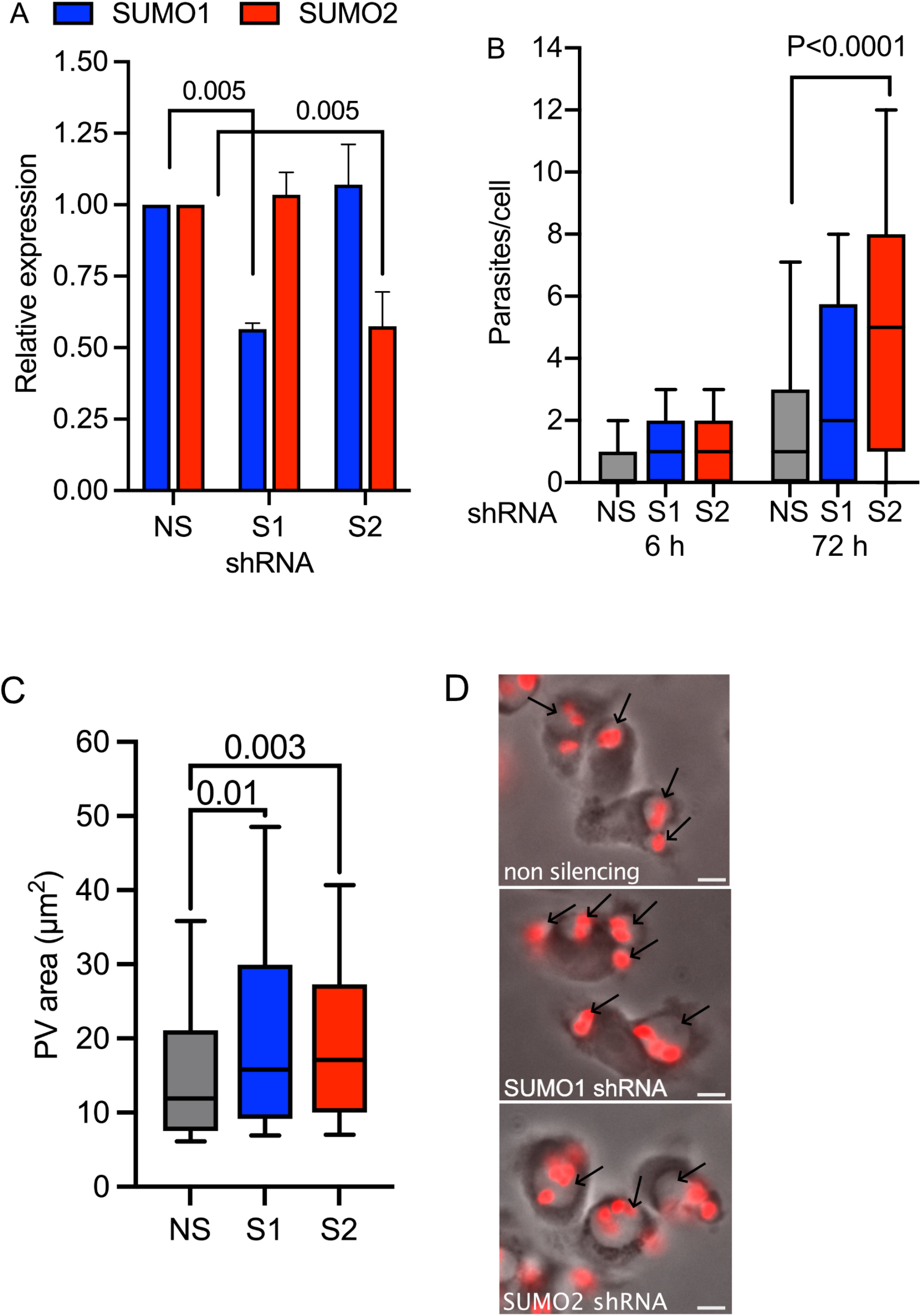

### SUMOylation inhibition promotes PV enlargement and drives expression of *ATP6V0D2*

Intracellular amastigotes from the *L. mexican*a group, including *L. amazonensis* induce the biogenesis of enlarged PVs, which favor their survival and replication. We investigated the hypothesis that reduction of SUMOylation could promote PV biogenesis. Indeed, macrophages with knockdown of SUMO1 or SUMO2 presented larger PVs than control cells (Figure 3C-D), consistent with the hypothesis that disruption of macrophage SUMOylation favors PV maturation and thereby enhances parasite growth.

The mechanisms driving the PV biogenesis are of great interest but still poorly understood. In our previous work we demonstrated that the scavenger receptor CD36 participates in the biogenesis of *L. amazonensis* PV while Pessoa *et al*. (2019) reported that an alternate subunit of the V-ATPase, Atp6v0d2, is involved in PV enlargement and parasite survival by regulating *CD36* expression and cholesterol retention [42]. Thus, we investigated if SUMOylation inhibition promotes the expression of *ATP6V0D2, CD36* or cholesterol retention.

THP-1 macrophages, control and SUMO-depleted, were infected with amastigotes and expression of *ATP6V0D2* and *CD36* determined by qRT-PCR. In wild type cells, parasite infection induced expression of *ATP6V0D2* 3 to 5-fold at 6 h to 24 h pi. Moreover, *ATP6V0D2* expression was significantly higher in SUMO1 and SUMO2 knockdown macrophages, including higher induction post infection (Fig. 4A). The expression of *CD36* was not consistently elevated by infection in control or knockdown cells, but SUMO1/2 knockdown cells presented generally higher levels (Fig. 4B).

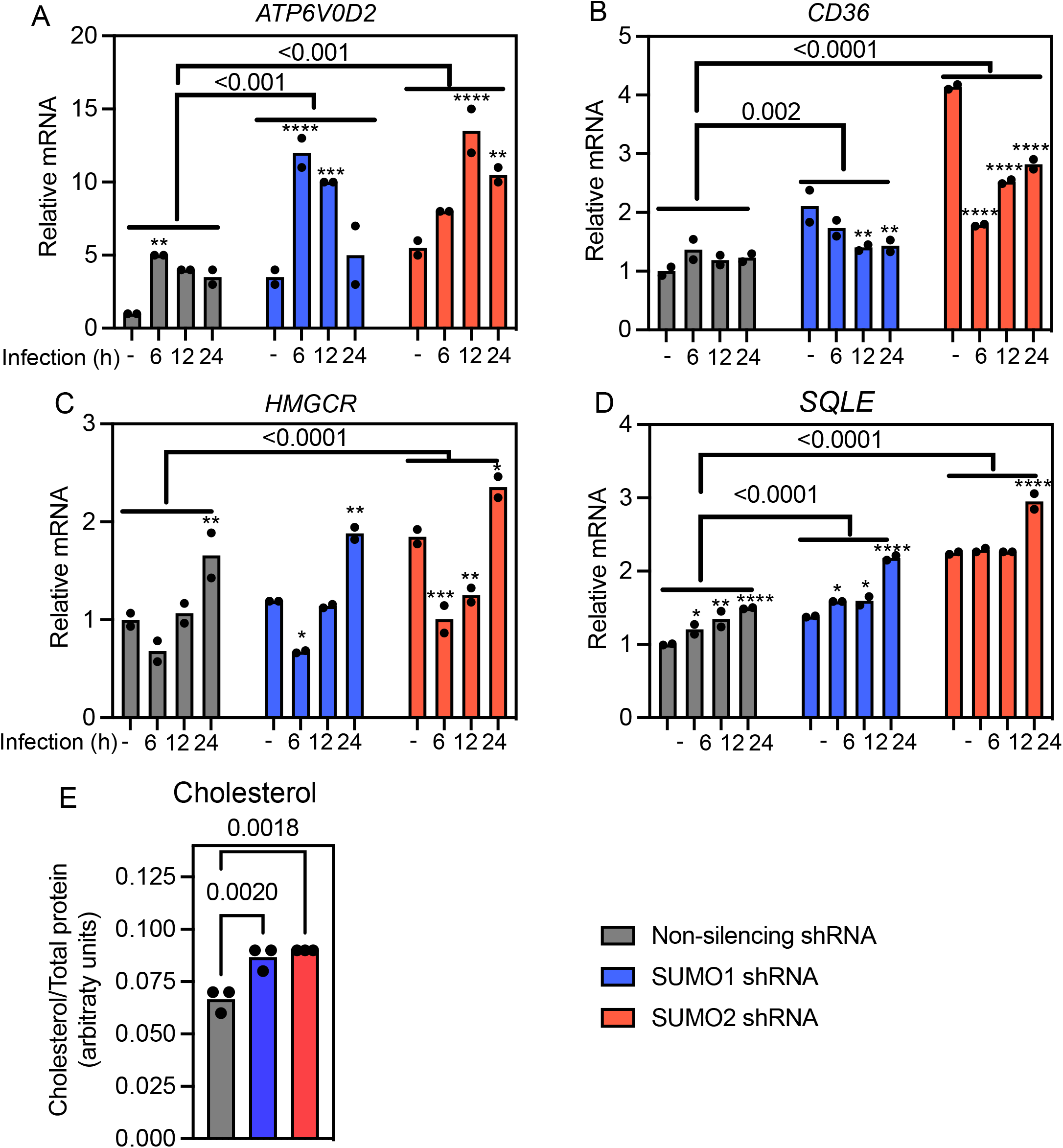

It was previously shown that knockdown of *Atp6v0d2* reduces the macrophage overall cholesterol levels and impairs PV biogenesis [42]. In fact, *SUMO2* depleted cells, with elevated *ATP6V0D2* expression, also presented higher levels two key genes of the cholesterol biosynthesis pathway, *HMGCR* and *SQLE*, while *SQLE* was similarly elevated only in *SUMO1* knockdown (Fig 4C-D). Moreover, lysates of SUMO depleted cells contained higher levels of total cholesterols (Figure 4E). Together, these results indicate that SUMOylation inhibition, perhaps through its effect on *ATP6V0D2* expression, increases expression of CD36 and some cholesterol biosynthetic genes and intracellular cholesterol levels, favoring PV biogenesis.

### *Atp6v0d2* expression is sufficient to increase free cholesterol, *CD36* expression and promote PV expansion in macrophages

Similar to human THP-1 macrophages, infection of RAW 264.7 cells with amastigotes promoted the expression of *Atp6v0d2* (Fig. 5A). To evaluate if *Atp6v0d2* induction is sufficient for the changes observed in SUMO depleted macrophages we generated RAW 264.7 macrophages stably expressing this gene. [Note, THP-1 cells were not used in these experiments because constitutive *ATP6V0D2* expression was toxic to THP-1 cells.] Interestingly, *ATP6V0D2* expressing RAW 264.7 cells phenocopied many aspects of SUMO depleted THP-1 macrophages, with increased the expression of CD36 and significantly higher levels of free cholesterols (Fig. 5B-C). However, *ATP6V0D2* overexpression did not induce the expression of cholesterol biosynthetic genes in RAW 264.7 cells (Figure S4), suggesting that higher levels of cholesterol observed in this cell line is linked to increased lipid intake by CD36. Importantly, *Atp6v0d2* expression strongly promoted PV biogenesis (Fig. 5D-E). Together, these results show that intracellular amastigotes inhibit SUMOylation to increase *ATP6V0D2* expression, which leads to increased cholesterol intake and drives PV enlargement and parasite proliferation, in human and mouse macrophages.

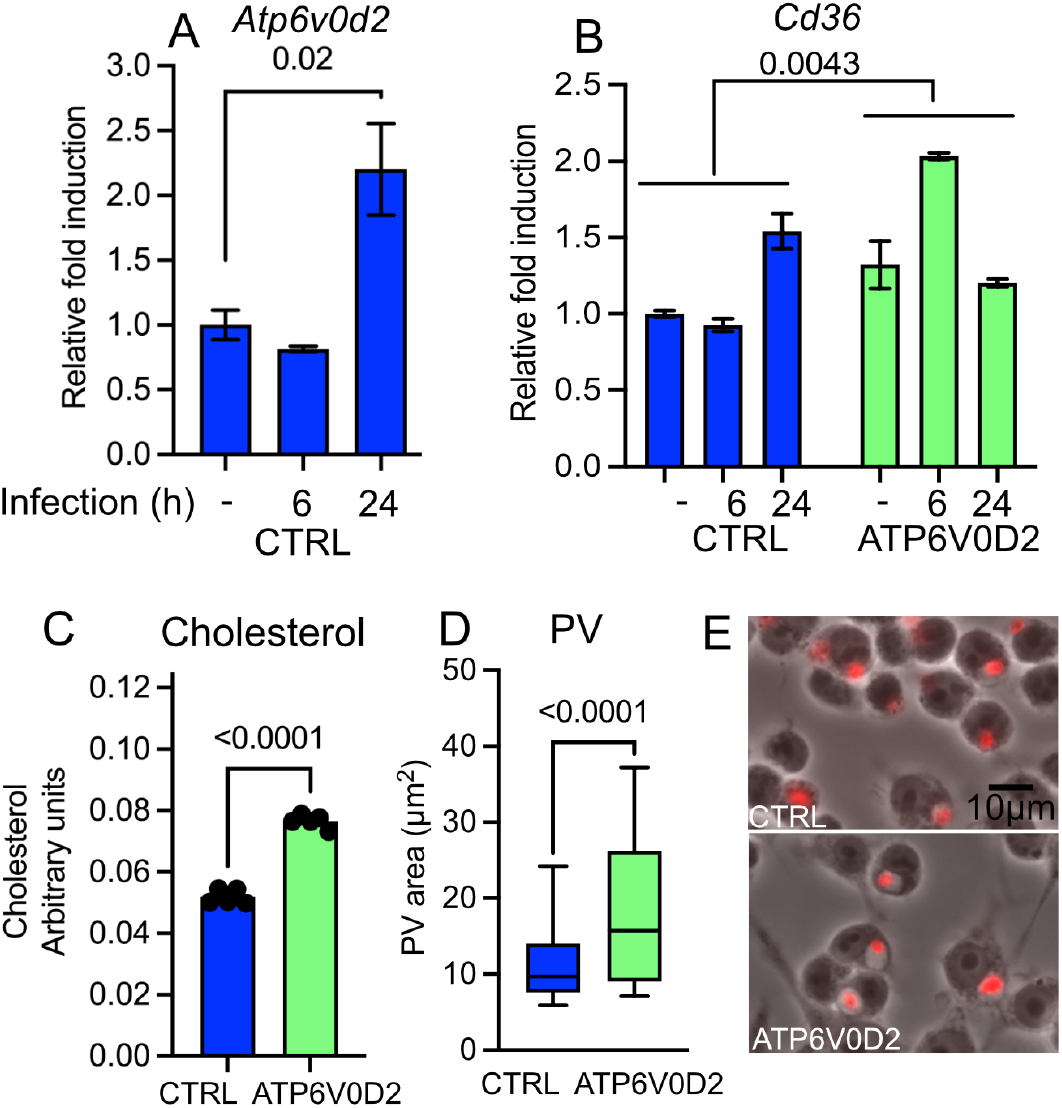

## Discussion

This work began with a genome-wide RNAi screen in *Drosophila* SL2 cells to identify novel factors involved in *L. amazonensis* amastigote infection. The screen was designed to identify factors involved in the initial infection steps, when phagocytes recognize and engulf parasites, potentially triggering cellular responses that can affect the downstream fate of infection such as PV biogenesis. *Drosophila* phagocytes have been successfully used to study several human pathogens, including intracellular bacteria, viruses, and protozoan parasites [26, 43-47]. The conservation of key recognition and signaling pathways involved in host defense and phagocytosis, including *Leishmania* uptake, was highlighted by the findings from our screen where 52 of 55 genes identified for their specific effects on *L. amazonensis* infection, are conserved between insects and humans.

We did not find common hits between our results and the published work from Peltan *et al*. (2012), who screened for *L. dononovani* amastigote infection factors also in *Drosophila* S2 cells [26]. This emphasizes the large phylogenetic distance between these two New World *Leishmania* species. Another explanation for the lack of concordane is that *L. donovani* screen used a dsRNA library with only 1,920 conserved genes compared to our genome wide library targeting 13,071 genes. The library used for *L. donovani* screen did not contain, for example, three SUMOylation factors, AOS1, Su(var)2-10 and UBA2, identified in our screen.

Based on the strong SUMOylation signature in the screen results, we explored the role of SUMOylation in *Leishmania* infection of mammalian macrophages. Interestingly amastigote, but not promastigote, infection quickly and potently inhibited host SUMOylation in both human and mouse macrophages, even inhibiting the induction of SUMO conjugation observed with heat shock. This drastic protein deSUMOylation seems to favor infection because knockdown of SUMOylation factors prior to infection caused higher proliferation of amastigotes within larger macrophages PVs. Together these data suggest *L. amazonensis* amastigotes actively interfere with the host cell SUMO machinery to enhance their own replicative niche.

SUMO regulates numerous cellular functions and activities, and human cells have 3,500 to 6,700 SUMOylated proteins, many of them SUMOylated on multiple residues [17, 48-50]. Not surprisingly, many pathogens sabotage this post-translational modification to facilitate infection. For example, bacteria *Salmonella typhimurium, Listeria monocytogenes, Shigella flexneri* and *Yersinia pestis* actively interfere the SUMOylation machinery during intracellular invasion [18-20, 51]. DeSUMOylation caused by *L. monocytogenes* inhibits TGF beta signaling, while *S. flexneri-induced* deSUMOylation increases permeability of the gut, favoring bacterial invasion and induces an exacerbated inflammatory response in mice [20]. Intracellular *S. typhimurium* reduces Rab7 SUMOylation and favors the formation of *Salmonella*-induced membrane filaments radiating from the Salmonella-containing vacuole which improves bacteria survival [21]. Finally, two protozoan parasites, which also live in parasitophorous vacuoles, *Plasmodium berghei* and *Toxoplasma gondii*, reduce protein SUMOylation to inhibit nuclear translocation of immune induced transcription factors, such as SMAD4, thereby reducing cellular responses to infection, resisting apoptosis, and favoring parasite infection [39].

Our data support the hypothesis that amastigotes cause a rapid inhibition of host protein SUMOylation to induce ATP6V0D2 expression and PV enlargement to facilitate survival of intracellular amastigotes. The enlarged PV, typical of parasites from the *L. mexicana* group, is a specialized organelle that protects the intracellular parasites from anti-parasitic host defense such as nitric oxide [52]. PV enlargement is actively driven by amastigotes, which manipulate macrophage vesicle trafficking and lipid metabolism/transport for this purpose; however the mechanisms used by *L. amazonensis* to regulate these processes are largely unknown [53, 54]. Here we propose that SUMOylation inhibition is a *Leishmania* virulence mechanism that increases availability of host cell cholesterol, promotes PV enlargement and favors infection.

One striking finding was that SUMOylation inhibition promotes the expression of *ATP6V0D2*, a protein previously reported to be involved in PV biogenesis [42]. Atp6v0d2 is best known as a component of vacuolar (V-)ATPases. V-ATPases are multiunit complexes composed by two domains: V_1_ domain, responsible for ATP hydrolysis and generation of energy for the proton translocation through the integral V_0_ domain. Atp6v0d subunits are part of the V_0_ domain and forms a connective stalk between subunits V_0_ and V1. While V-ATPases are present in all cells, *Atp6v0d2* is expressed in specific tissues (renal, epididymis, osteoclast, dendritic cells, macrophages), while the more common d-subunit, Atp6v0d1 is expressed ubiquitously [55, 56]. More recently, Atp6v0d2 has been shown to participate in several cellular processes distinct from V-ATPase activity, including fusion of osteoblasts [57], autophagosome-lysosome fusion and inflammasome activation [58], and the degradation of transcription factors [56]. Thus, this alternate V-ATPase subunit appears to have roles both in vacuolar acidification and in unrelated processes that often involve membrane fusion.

This study links *ATP6V0D2* expression to parasite-triggered deSUMOylation. Previous work has linked Atp6v0d2, Cd36 and cholesterol levels [42]. The role of Cd36 in PV biogenesis was previously reported by our group, where the receptor favors fusion of endolysosomes to PV [25]. Since we observed that *ATP6VOD2* (over)expression had variable effects on cholesterol biosynthetic genes, in mouse or human macrophages but consistent effects on *CD36* expression, we favor the idea that the increased uptake of cholesterol observed in all these models is primarily mediated by elevated CD36 uptake. Interestingly, our previous work showed CD36 localized to PV membrane in contact with parasites, suggesting this lipid transporter may directly delivery cholesterol to PV membrane, lumen and/or to the parasite membrane. Through these modulations of *ATP6V0D2* expression and cholesterol levels, amastigote-triggered deSUMOylation has profound effects parasite intracellular growth.

One unanswered question is how SUMOylation inhibition promotes *ATP6V0D2* expression. The role of SUMOylation in the control of gene expression is well-established, particularly via the SUMOylation of transcription factors [59]. Although the effect of transcription factor SUMOylation varies, in most cases it reduces transcription of target genes [60]. Interestingly, a publicly available ChIP-seq dataset from mouse dendritic cells shows that a 588 bp DNA region, 32kb upstream of *Atp6v0d2*, is significantly enriched in SUMO2-conjugated chromatin [61]. Therefore, one possibility is that amastigote-induced deSUMOylation of transcription factor(s), which bind near this element near *Atp6v0d2*, drive higher expression of *Atp6v0d2*. Some potential candidates are the transcription factors CTCF, RAD21, Zinc Finger protein 143, c-FOS and PPARγ, whose activities are known to be modulated by SUMOylation and are linked in *Atp6v0d2* expression [62, 63].

In summary, our *Drosophila* screen has proven to be effective for the identification of novel host factors involved in *Leishmania-host* cell interactions. Our results identify a novel *L. amazonensis* virulence strategy, whereby amastigotes drastically reduce host cell protein SUMOylation to increase *ATP6V0D2* expression and cholesterol availability, as a strategy to promote their own proliferation. The *Leishmania* parasitophorous vacuole has clear parallels to the replicative niche of other intracellular bacterial pathogens and it will be interesting to learn, from future studies, if similar processes are involved. Further in-depth studies focused on the mechanistic interactions between protein SUMOylation and parasite growth, as well as on the other hits found in the *Drosophila* screen may reveal new targets for leishmaniasis treatment.

## Materials and Methods

### Ethics statement

Our lab conducted all experiments according to the guidelines of the American Association for Laboratory Animal Science and approved by the Institutional Animal Care and Use Committee at the University of Massachusetts Medical School (Docket#: A-2056-19).

### Mice

C57BL/6 and Balb/c mice were obtained from The Jackson Laboratory. Seven to twelve-week-old mice were maintained under pathogen-free conditions at the University of Massachusetts Medical School animal facilities.

### Macrophages

BMDM: femurs and tibia were dissected from mice and the bone marrow was flushed with PBS using a 30G needle connected to a 10 ml syringe. The cells were cultivated in RPMI supplemented with 30% L929 cell-conditioned medium and 20% FBS in bacteriological petri dishes (4×10^6^ cells/plate) at 37 °C and 5% CO_2_ [67]. The cultures were re-fed on day 3 and used on day 7-10. THP-1 monocytes were differentiated to macrophages by stimulation with 100 ng/mL of phorbol 12-myristate 13-acetate (PMA) in RPMI 10% FBS for 2 days at 37°C, washed and cultured for more 24 h in complete media without PMA, prior use. RAW 264.7 cells were cultivated in DMEM supplemented with 10% FBS.

### Macrophage transduction

SUMO1 and SUMO2 were knocked down using GIPZ lentiviral particles from GE Dharmacon V3LHS_404077 and V3LHS_412780. ATP6V0D2 (NM_152565.1) was cloned with a C-terminus tag in the lentiviral vector PCX4-PURO and used to generate lentiviral particles and transduce macrophages. Cells were selected with puromycin before use.

### Confocal microscopy

SL2 cells were seeded (2×10^5^ cells per dish) in 35mm dish with glass bottom one day before infection. Cells were infected with amastigotes and at determined times, cell membranes were stained with CellMask deepRed for 10 min, and acidic organelles were stained with Lysotracker Blue DND-22 (Invitrogen). Confocal images were taken using a Leica SP8 confocal microscopy using a 63x objective. Images were edited using Fiji software [64].

### *Leishmania* parasites

GFP-transfected *L. amazonensis* (MHOM/BR/1973/M2269), and dsRed-transfected (RAT/BA/LV78) [65] were generously donated by Dr. Silvia R. Uliana (ICB-USP, Brazil) and Dr. Kwang-Poo Chang (Rosalind Franklin University of Medicine and Science), respectively. Promastigotes were cultivated *in vitro* in M199 medium supplemented with 10% FBS and 30 μg/mL Hygromycin B (GFP transfected) or 5 μg/mL of Tunicamycin (dsRed transfected) at 26 °C.

### Production of amastigote forms

Promastigote forms were differentiated axenically to amastigote by cultivating the parasites with 199 media supplemented with 0.25% glucose, 0.5% trypticase, 40 mM sodium succinate (pH 5.4) and 20% FBS, at 1×10^7 cells/mL at 34 °C for 3 days. Axenic amastigotes were then used to infect Balb/c or C57BL/6 BMDM cultivated in 175 cm^2^ T-flasks (3×10^7^ cells/flask) at a multiplicity of infection of 3 parasites/BMDM. At least 4 days after infection cells were harvested from T-flasks using a cells scraper and disrupted in PBS at 4°C using a glass dounce tissue grinder with teflon rod. Parasites were purified by two-step centrifugations, one at 210*g* for 8 min to remove intact BMDMs and cell debris, and then supernatant was centrifuged at 675*g* for 12 min to precipitate amastigotes.

### HTS screening

For genome wide screen a library of 13,071 dsRNA against *D. melanogaster* genes (Ambion) distributed in 32 384-well plates with clear bottom (Costar) was utilized. SL2 cells were maintained in complete media (Schneider media (Gibco), 10% fetal bovine serum (Optima, Atlanta Biologicals), 100 iU of penicillin and streptomwycin) at 27° C. After changing to serum-free media, 20,000 cells in ten μL of was added to each well, which contained 250 ng dsRNA, and incubated for 1 h, then 20 μl of complete media was added per well and incubated for 72 h for knockdown. GFP expressing amastigotes harvested from BMDMs (10^4^ in 5μl of Schneider media per well) were added to each well and the plates centrifuged at 300 x G for 5 min and kept at 27C for 90 min. The media was carefully removed from wells and cells were fixed with 20μl of 3.2% formaldehyde (EM grade, EMS) for 30 min at room temperature and not permeabilized.

The cultures were washed once with PBS and blocked with 1% bovine serum albumin (BSA) for 1 h at 37° C. Extracellular parasites were immunostained with serum of an infected Balb/c diluted 1:4000 in PBS with 1% BSA, and a secondary goat anti-mouse conjugated to Alexafluor 594 1:1000 (Invitrogen). The DNA of cells were counterstained using 0.1μg/ml Hoechst 33342 added to the secondary antibody solution, the general scheme for the method is shown in Figure S2. Three fields per well, close to the center of wells, were imaged using a 20x objective using an ImageXpress Micro automated microscope and analyzed using MetaXpress software (Molecular Devices). Number of SL2 cells (Hoechst 33342 staining, total parasites (GFP positive), and extracellular parasites (GFP and AlexaFluor 594 positive), intracellular parasites (GFP positive and AlexaFluor 594 negative) were scored and averaged for the three images of each well. The infection rate (number of intracellular parasites per SL2 cells) was log-transformed, and the median and interquartile range were used to calculate a Z-score: (log10(%infection)-log10(median))/(IQR*0.74) for each plate [30]. The entire screen was performed in duplicate and wells with Robust Z-scores of infection ≥ 1.5 or ≤ −2.0 in both replicates were considered ‘hits’. Knockdowns that reduced the SL2 numbers by −2 Z-scores received lower priority to further studies.

### dsRNA Synthesis and secondary screen

Primer sequences to generate dsRNA sequences with no off-targets were obtained from DRSC/TRiP Functional Genomics Resources (www.flyrnai.org) or manually designed using SnapDragon (https://www.flyrnai.org/cgi-bin/RNAi_find_primers.pl) (Supplemental Table 2). After PCR using genomic DNA from flies, dsRNA was synthesized using MEGAscript T7 transcription kit (Invitrogen) following manufacturer recommendations.

The secondary screen plates were arrayed with dsRNA (250 ng) using on-site synthesized dsRNA free of off targets. Genes were considered hits when two of the triplicates Robust Z-scores in infection was ≥1.5 or ≤1.5 using dsRNA against 180 wells treated with dsRNA against β-galactosidase.

### Phagocytosis of *E. coli* and *S. aureus*

SL2 cells were soaked with 1.25 μg of dsRNAs in 96 well plate for 3 days prior to phagocytosis assays. Fluorescein-labeled *E. coli* (K-12 strain), and *Staphylococcus aureus* (Wood strain, without protein A) (Molecular Probes), were washed 3 times with PBS by centrifugation and sonicated for 3 times at 50 KHz for 20 s. Twenty micrograms of bacterial particles were added to each well at 4°C, the plates were centrifuged 300xg for 3 min at 4 °C, and incubated in a 27° C water bath for 20 or 40 min (for *E. coli* and *S. aureus*, respectively). On ice bath, cells and bacteria were removed from plates by pipetting 300 ul of a 0.1% Trypan Blue solution in PBS pH 5.4 to quench the fluorescein from extracellular bacteria. The bacteria inside live cells was quantified by flow cytometry [29].

### Bioinformatic analysis

The clustering and enrichment analysis of the screen hits was performed in PANTHER Classification System [66].

### Western Blot

Macrophages were lysed in standard lysis buffer containing 10% glycerol, 1% Triton X-100, 20 mM Tris, 150 mM NaCl, 24 mM β-glycerol phosphate, 2 mM EDTA, 1 mM 1,4 dithiothreitol (DTT), 1 mMsodium orthovanadate, 5 mM N-ethylmaleimide, 0.5 U/μl Benzonase, 1× protease inhibitor mixture (Halt, Thermofisher). After protein quantification by Bradford protein assay (Bio-Rad), Laemmli sample buffer 4x was added to samples and heated for 3 min at 95°C. For whole cell lysate analysis, 50 μg of total protein was separated by SDS-PAGE using 8% or 4-20% polyacrylamide gels, and transferred to PVDF membranes at 350 mA using BioRad Mini Protean and Transblot system (Bio-Rad). Membranes were probed with antibodies against SUMO1 or SUMO2/3 (Cell Signaling C9H1 and 18H8) using manufacturer conditions. For loading controls, the same blots were probed with anti-actin or anti-GAPDH, and then stained with Coomassie blue (sc8432, Santa Cruz, G9295 Millipore Sigma).

### Production of retroviral vectors and transduction of immortalized mouse macrophages

The full length mouse SUMO2 fused to HA (N-terminus) from a plasmid (Addgene 48967) [67] were inserted in the retroviral transfer plasmid pCX4 Puro. For the viral production, HEK 293T cells were transfected with the pCX4 HA-SUMO2, MLV gag-pol, and VSVG using GeneJuice transfection reagent (EMD Millipore) following the manufacturer’s recommendations. Media was replaced at 24 h and collected 48 h after transfection, filtered through a 0.45 μm pore filter and stored at −80 °C until use. Virus for shRNA transduction were generated using shRNA library (GIPZ, Dharmacon): SUMO2 (V3LHS_412780), SUMO1-(V3LHS_404077), SAE1 V3LHS_304633, UBE2l (V3LHS_376933). THP-1 cells were cultured in the presence of virus particles and 8 μg/ml Polybrene for 24 h, and then the transduced cells were selected with 3 μg/ml of puromycin for 4 days. Cells were frozen and when the thawed aliquots were subcultured for up to 3 weeks.

### ATP6V0D2 Plasmid

Human ATP6V0D2-FLAG NM_152565.1 was obtained from Sino Biological (HG23801-CF) and cloned in PCX4-Puro lentiviral plasmid.

### Macrophage infection and PV area measurement

THP-1 macrophages were infected with amastigotes for 2 h and the coverslips were washed 3 times with RPMI media and transferred to a new plate. One day after infection the cultures were fixed with 4% paraformaldehyde for 15 min and mounted over a glass slide for microscopic imaging in a fluorescence microscope. The area of PV was manually measured using an elliptical tool in FIJI.

### RNA and qRT-PCR

Total RNA from macrophages were isolated using the TRIzol reagent (Invitrogen) following the manufacturer’s recommendations. cDNA was synthesized using iScript gDNA Clear cDNA synthesis kit (BioRad) and quantitative PCR analysis was performed using SYBR Green (BioRad). The specificity of amplification was assessed for each sample by melting curve analysis and relative quantification was performed using a standard curve with dilutions of a standard.

Oligonucleotides used in qPCR reactions:

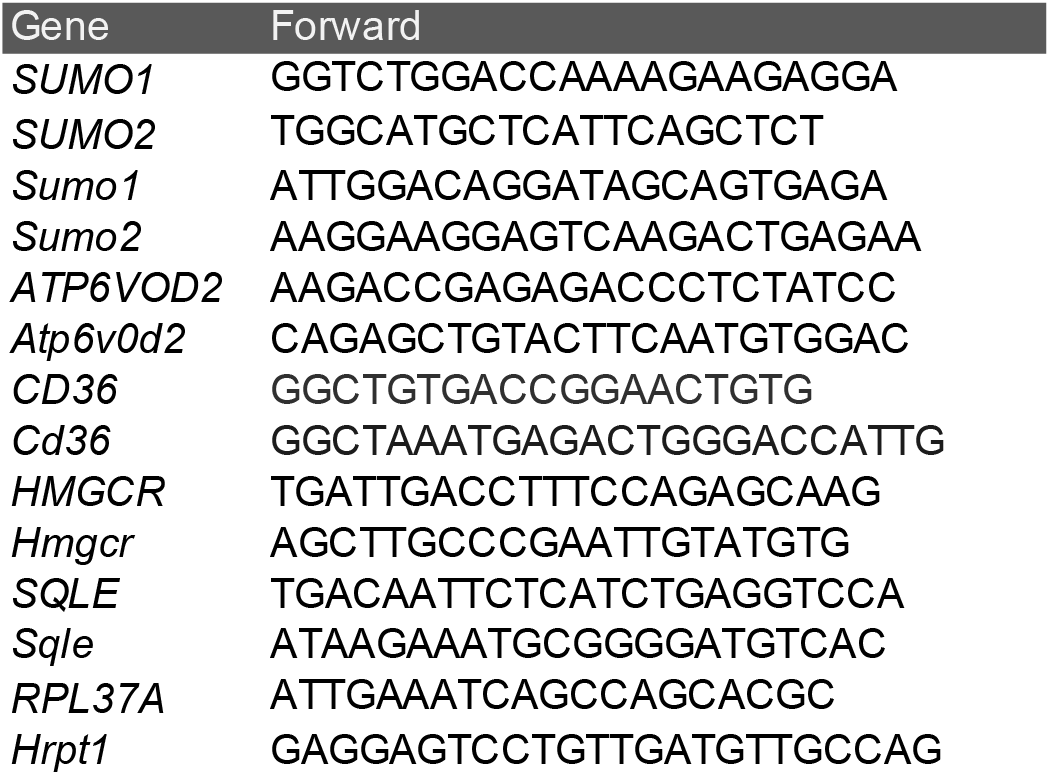

### Cholesterol measurement

Macrophages were seeded in 96 well plates (5×10^4 cells per well) and one day later, free and total cholesterol concentration in lysates was determined using Cholesterol/Cholesterol Ester-Glo Assay (Promega) following manufacturer recommendations. Cholesterol concentrations were normalized by the protein concentration determined by Bio-Rad Protein Assay Dye (Bio-Rad).

### Statistical analysis

The statistical analyses were performed using GraphPad Prism version 6.00 for Mac OS X, GraphPad Software, La Jolla California USA, www.graphpad.com. The tests and the criteria used for each comparison are reported in the Figure legends.

## Notes

### Competing Interest Statement

The authors have declared no competing interest.

