## Supplementary figures and images for "*Leishmania amazonensis* sabotages host cell SUMOylation for intracellular survival"

### Supplemental Figure 1

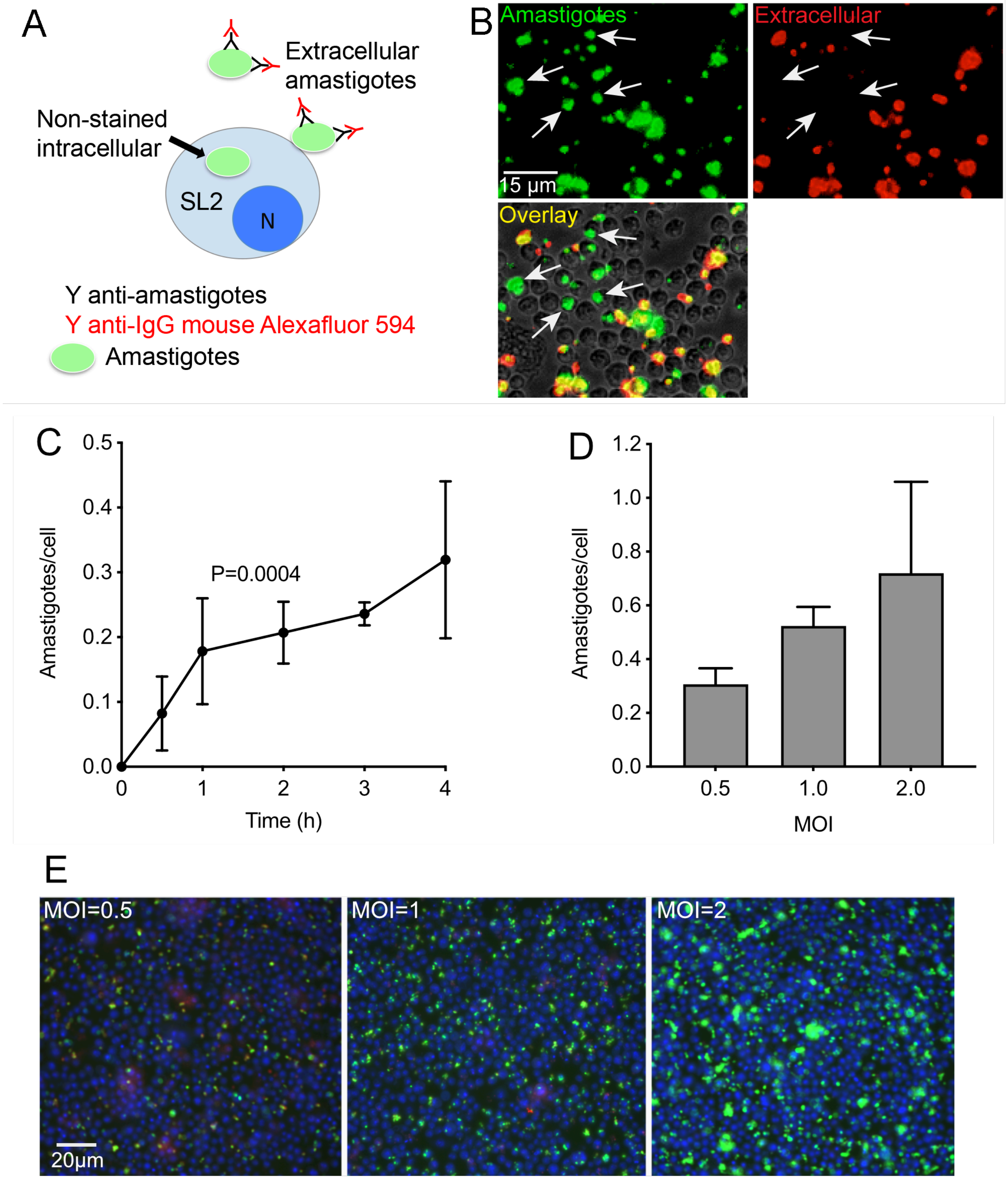

### Supplemental Figure 2

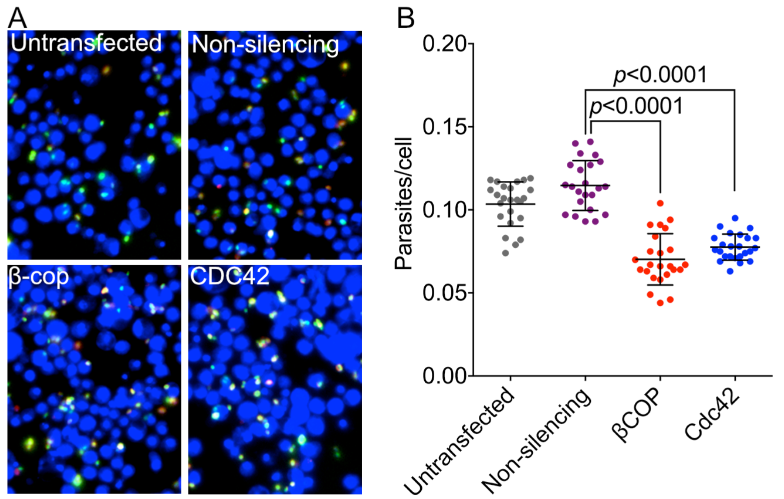

### Supplemental Figure 3

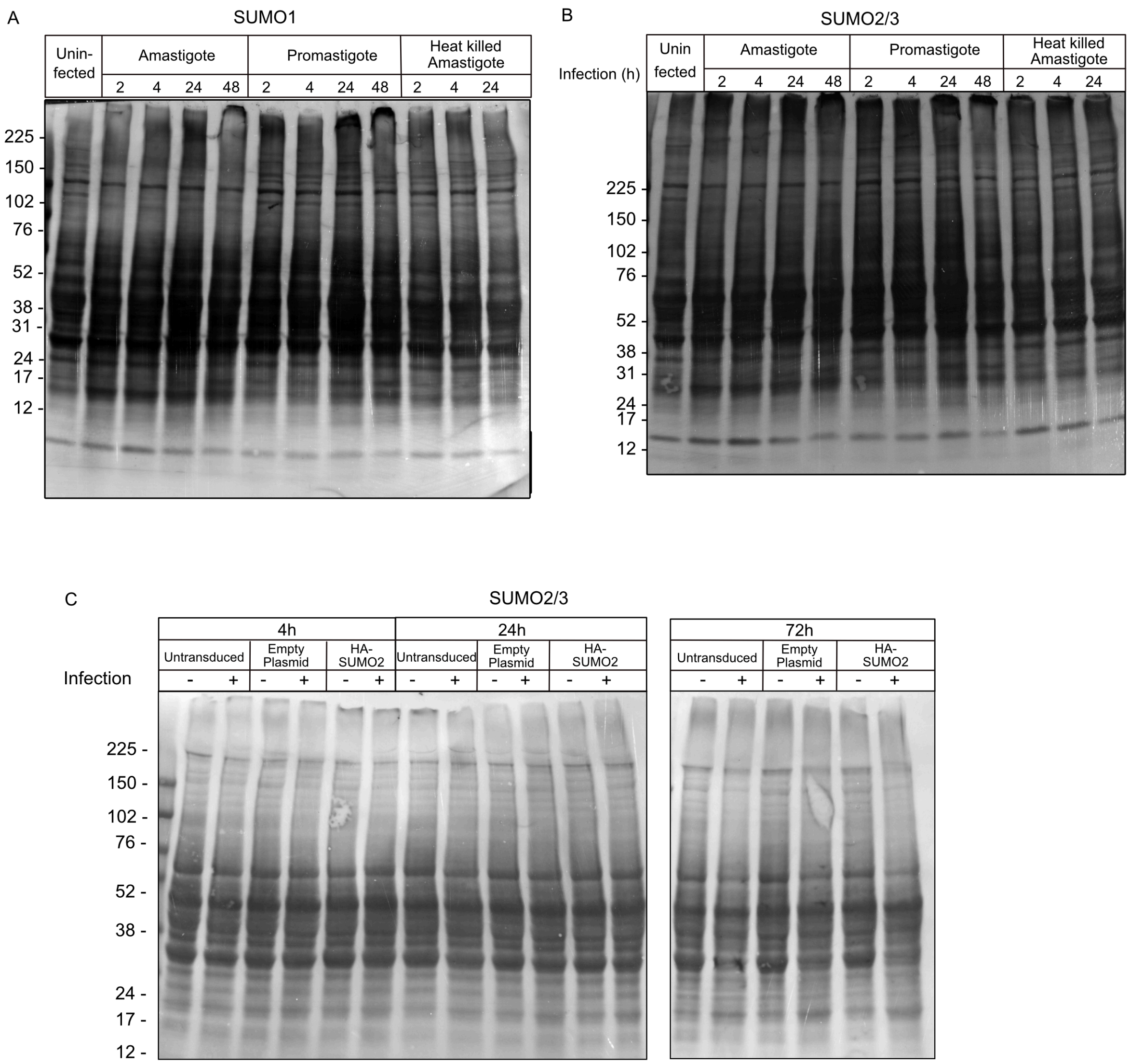

### Supplemental Figure 4

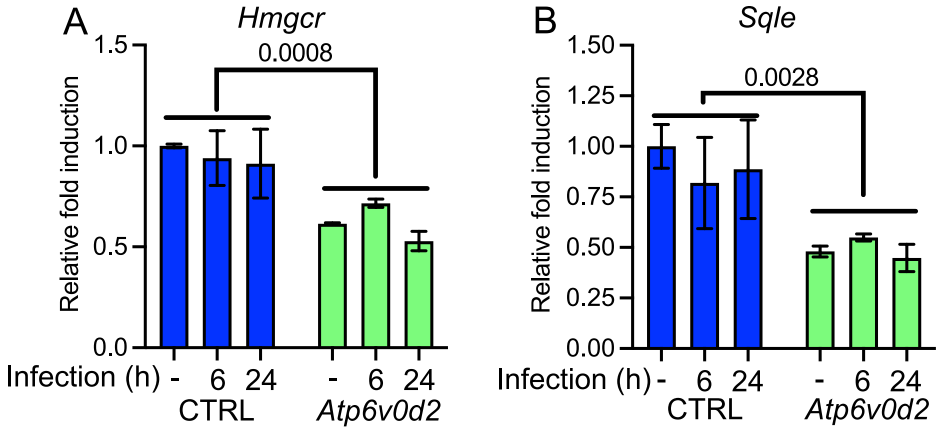
