## Supplemental Table 1 for "*Leishmania amazonensis* sabotages host cell SUMOylation for intracellular survival"

| | | | | | | | | | | | | | | | | | | | |
|---|---|---|---|---|---|---|---|---|---|---|---|---|---|---|---|---|---|---|---|
| | Flybase id | | | 1st screen | | | | Validation Screen | | | | | | Relative Bacterial burden | | | | | |
| | | target_gene | symbol | Robust Z-score Infection | | Robust Z-score SL2 | | Robust Z-score Infection | | | Robust Z-score SL2 | | | *E. coli* | | | *S. aureus* | | |
| | | | | Set 1 | Set 2 | Set1 | Set 2 | Set1 | Set 2 | Set3 | Set1 | Set 2 | Set3 | Set1 | Set2 | Set3 | Set1 | Set2 | Set3 |
| Leishmania curated infection hits | | | | | | | | | | | | | | | | | | | |
| 1 | FBgn0027594 | CG2086 | drpr | -7.6 | -2.3 | -1.1 | 1.4 | -4.5 | -3.9 | -5.0 | 1.2 | 1.1 | 0.5 | 1.2 | 1.2 | 1.1 | 1.0 | 0.9 | 1.0 |
| 2 | FBgn0001297 | CG15509 | kay | -2.9 | -2.9 | -0.3 | -2.8 | -4.3 | -3.4 | -4.9 | -1.6 | 0.6 | -2.2 | 1.2 | 1.0 | 1.1 | 0.7 | 0.4 | 0.8 |
| 3 | FBgn0040066 | CG17437 | wds | -2.2 | -2.2 | -1.1 | -1.4 | -3.8 | -4.5 | -3.9 | 1.1 | 0.9 | 1.3 | 1.0 | 1.0 | 0.9 | 0.7 | 0.8 | 0.9 |
| 4 | FBgn0028539 | CG31731 | CG31731 | -6.0 | -5.1 | 0.4 | -0.9 | -3.3 | -4.1 | -4.3 | 0.8 | 1.1 | -0.6 | 1.1 | 1.3 | 1.2 | 0.9 | 0.9 | 1.0 |
| 5 | FBgn0016696 | CG4268 | Pitslre | -2.6 | -2.4 | -5.4 | -5.3 | -5.1 | -3.3 | -3.1 | -3.8 | -1.3 | -2.6 | 1.0 | 1.0 | 1.1 | 0.6 | 0.7 | 0.8 |
| 6 | FBgn0086357 | CG9539 | Sec61alpha | -2.0 | -2.8 | -1.2 | -1.1 | -4.7 | -3.3 | -3.3 | -1.0 | 0.8 | -0.3 | 1.2 | 0.8 | 0.7 | 0.6 | 0.7 | 0.7 |
| 7 | FBgn0037573 | CG7483 | eIF4AIII | -6.1 | -6.5 | -1.9 | -3.4 | -4.1 | -4.2 | -2.8 | -0.5 | -0.9 | -0.3 | 1.2 | 1.1 | 1.2 | 1.0 | 1.0 | 1.1 |
| 8 | FBgn0002736 | CG9401 | mago | -6.2 | -2.7 | 0.6 | -1.8 | -3.4 | -2.3 | -5.3 | -0.5 | 0.2 | -1.1 | 1.0 | 1.0 | 0.9 | 0.7 | 0.7 | 0.7 |
| 9 | FBgn0000559 | CG2238 | Ef2b | -4.6 | -4.8 | -1.7 | -3.0 | -5.3 | -2.7 | -2.9 | -2.0 | -1.0 | -2.5 | 1.5 | 1.4 | 1.3 | 1.2 | 1.1 | 1.1 |
| 10 | FBgn0038473 | CG3983 | Ns1 | -4.1 | -2.2 | -1.7 | -0.9 | -3.3 | -3.9 | -3.0 | 0.3 | 1.3 | -0.7 | 1.4 | 1.7 | 1.4 | 1.1 | 1.2 | 0.9 |
| 11 | FBgn0028473 | CG8801 | Non1 | -4.7 | -3.8 | -1.0 | -2.0 | -2.5 | -3.0 | -4.4 | -2.3 | 1.9 | 0.9 | 1.4 | 1.7 | 1.3 | 1.3 | 1.1 | 1.2 |
| 12 | FBgn0037899 | CG6764 | RpL24-like | -5.2 | -3.1 | -4.8 | -1.0 | -5.1 | -3.1 | -1.7 | 1.5 | 1.6 | 2.2 | 1.3 | 1.4 | 1.3 | 1.5 | 1.2 | 1.4 |
| 13 | FBgn0030007 | CG2263 | α-PheRS | -2.3 | -2.4 | 0.1 | -2.8 | -3.6 | -3.2 | -2.7 | -1.3 | 0.6 | 0.0 | 1.3 | 1.5 | 1.4 | 1.0 | 1.0 | 1.6 |
| 14 | FBgn0027553 | CG32721 | NELF-B | -4.6 | -3.3 | -3.6 | -0.2 | -3.1 | -2.7 | -3.5 | -0.9 | 0.2 | 0.3 | 1.2 | 1.2 | 1.2 | 0.8 | 0.7 | 1.0 |
| 15 | FBgn0025366 | CG5277 | Ip259 | -3.6 | -3.9 | -1.6 | -2.6 | -3.5 | -2.4 | -2.9 | 0.7 | 0.6 | 0.7 | 1.6 | 1.5 | 1.6 | 1.7 | 1.5 | 1.2 |
| 16 | FBgn0037371 | CG2097 | Sym | -3.6 | -3.0 | 0.7 | -1.5 | -4.7 | -1.8 | -2.2 | -1.1 | 1.3 | -0.5 | 1.3 | 1.3 | 1.1 | 0.8 | 0.9 | 1.2 |
| 17 | FBgn0033661 | CG13185 | CG13185 | -4.6 | -2.7 | -0.4 | -1.0 | -3.6 | -2.6 | -2.4 | 0.1 | 1.3 | 2.3 | 1.2 | 1.1 | 1.1 | 0.8 | 0.6 | 1.0 |
| 18 | FBgn0023213 | CG10811 | eIF4G1 | -3.9 | -4.7 | 0.2 | -4.1 | -2.1 | -2.5 | -3.9 | -2.9 | -0.6 | -3.0 | 1.6 | 1.8 | 1.4 | 1.0 | 0.9 | 1.3 |
| 19 | FBgn0267376 | CG6584 | SelR | -2.3 | -2.1 | -2.8 | -0.1 | -3.0 | -1.9 | -3.4 | -6.4 | -4.8 | -6.0 | 1.1 | 1.1 | 1.1 | 0.9 | 0.8 | 1.0 |
| 20 | FBgn0032453 | CG6180 | CG6180 | -3.6 | -2.8 | -0.8 | -2.2 | -2.1 | -3.5 | -2.6 | -0.9 | -2.1 | -1.1 | 1.4 | 1.3 | 1.4 | 1.1 | 1.1 | 1.7 |
| 21 | FBgn0261609 | CG9946 | eIF-2alpha | -3.2 | -5.8 | -0.6 | -3.4 | -2.2 | -2.3 | -3.4 | -5.6 | -2.1 | -3.1 | 1.7 | 1.6 | 1.8 | 1.9 | 1.8 | 1.6 |
| 22 | FBgn0002791 | CG3060 | mr | -4.4 | -2.3 | -1.4 | -2.6 | -3.7 | -2.0 | -2.0 | -3.5 | -0.9 | -2.1 | 0.9 | 0.7 | 0.5 | 0.4 | 0.5 | 0.4 |
| 23 | FBgn0259791 | CG6897 | bora | -2.2 | -2.3 | -2.7 | 0.3 | -1.9 | -3.4 | -2.2 | -0.2 | -0.3 | -0.5 | 1.0 | 1.0 | 0.9 | 0.7 | 0.5 | 1.0 |
| 24 | FBgn0015834 | CG8882 | Trip1 | -7.5 | -5.2 | -1.6 | -3.9 | -3.6 | -2.0 | -2.0 | -7.1 | -3.9 | -5.1 | 1.7 | 1.5 | 1.6 | 1.5 | 1.4 | 1.5 |
| 25 | FBgn0047114 | CG31142 | CG31142 | -2.2 | -2.6 | 0.1 | -1.0 | -3.5 | -2.4 | -1.7 | 0.4 | 0.7 | -1.3 | 1.2 | 1.3 | 1.1 | 1.0 | 0.8 | 1.3 |
| 26 | FBgn0266673 | CG6159 | sec10 | -2.3 | -4.4 | -0.7 | 0.1 | -2.1 | -2.8 | -2.2 | -0.5 | -0.8 | -1.0 | 1.1 | 1.3 | 1.2 | 1.0 | 0.8 | 1.2 |
| 27 | FBgn0029512 | CG12276 | Aos1 | -3.2 | -2.1 | -1.9 | -0.8 | -1.6 | -2.5 | -2.5 | -1.8 | -0.9 | -1.6 | 1.2 | 1.2 | 1.1 | 0.9 | 0.7 | 1.1 |
| 28 | FBgn0039233 | CG7006 | CG7006 | -2.1 | -2.7 | 0.9 | -0.7 | -2.9 | -1.8 | -1.6 | 2.5 | 1.3 | 1.5 | 1.2 | 1.3 | 1.1 | 1.0 | 0.9 | 1.1 |
| 29 | FBgn0034231 | CG11419 | APC10 | -2.9 | -4.8 | -1.8 | -3.4 | -1.7 | -2.6 | -1.8 | -1.4 | -0.2 | -0.1 | 0.8 | 1.0 | 0.9 | 0.8 | 0.7 | 0.7 |
| 30 | FBgn0260456 | CG4806 | CG4806 | -3.6 | -5.1 | -1.6 | -2.3 | -1.8 | -1.7 | -1.8 | -0.2 | 1.1 | 0.4 | 1.3 | 1.3 | 1.2 | 0.9 | 0.9 | 1.1 |
| 31 | FBgn0035524 | CG32253 | CG32253 | -8.0 | -2.5 | -5.5 | -1.3 | -3.7 | -3.9 | -1.1 | 0.3 | 1.4 | 0.8 | 0.9 | 1.5 | 1.3 | 1.2 | 1.4 | 1.4 |
| 32 | FBgn0010356 | CG7704 | Taf5 | -6.2 | -2.8 | -1.7 | -3.9 | -4.8 | -2.7 | -1.3 | -5.9 | -2.1 | -3.0 | 1.0 | 0.8 | 1.1 | 0.7 | 0.8 | 1.2 |
| 33 | FBgn0022213 | CG13281 | Cas | -4.1 | -4.4 | -1.7 | -4.9 | -4.3 | -3.2 | -1.3 | -6.8 | -3.7 | -4.4 | 1.5 | 1.4 | 1.6 | 0.8 | 0.7 | 0.9 |
| 34 | FBgn0266916 | CG31258 | Cenp-C | -3.0 | -4.3 | -1.0 | -4.5 | -4.7 | -2.4 | -1.4 | -5.1 | -2.2 | -2.4 | 1.1 | 1.0 | 1.1 | 0.6 | 0.6 | 0.7 |
| 35 | FBgn0035422 | CG12740 | RpL28 | -6.2 | -4.6 | -1.7 | -3.8 | -4.2 | -3.5 | -0.9 | -6.3 | -4.4 | -4.3 | 1.5 | 1.4 | 1.6 | 1.1 | 1.1 | 1.3 |
| 36 | FBgn0039302 | CG11856 | Nup358 | -2.5 | -5.4 | -1.2 | -6.5 | -1.1 | -2.6 | -4.1 | -8.1 | -5.3 | -6.3 | 1.0 | 0.9 | 0.9 | 0.3 | 0.2 | 0.4 |
| 37 | FBgn0037550 | CG9667 | CG9667 | -2.4 | -4.1 | -1.2 | -5.0 | -1.0 | -3.3 | -3.2 | -8.5 | -4.9 | -6.0 | 1.4 | 1.3 | 1.3 | 1.0 | 1.0 | 1.0 |
| 38 | FBgn0261396 | CG42641 | Dox-A2 | -3.2 | -2.7 | -0.3 | -6.0 | -2.7 | -1.2 | -3.5 | -8.7 | -5.1 | -5.6 | 1.0 | 0.8 | 0.9 | 0.5 | 0.5 | 0.6 |
| 39 | FBgn0035900 | CG6694 | ZC3H3 | -3.2 | -3.6 | -1.7 | -5.6 | -3.9 | -0.9 | -2.4 | -10.7 | -6.6 | -6.5 | 1.2 | 0.6 | 0.8 | 0.3 | 0.4 | 0.6 |
| 40 | FBgn0027088 | CG6778 | Aats-gly | -4.2 | -2.8 | -1.5 | -2.7 | -2.9 | -2.5 | -1.4 | -1.0 | 0.2 | 0.3 | 1.1 | 1.0 | 1.2 | 1.4 | 1.2 | 1.0 |
| 41 | FBgn0038320 | CG4931 | Sra-1 | -2.9 | -4.7 | 0.0 | -0.6 | -0.9 | -2.4 | -3.2 | -2.6 | -0.5 | -2.0 | 0.9 | 0.7 | 0.7 | 0.5 | 0.4 | 0.6 |
| 42 | FBgn0261602 | CG1263 | RpL8 | -4.2 | -3.0 | -0.5 | -4.7 | -2.7 | -2.1 | -1.1 | -9.3 | -5.3 | -5.2 | 1.7 | 1.8 | 1.7 | 1.3 | 1.1 | 1.4 |
| 43 | FBgn0036828 | CG6841 | CG6841 | -3.3 | -3.7 | -1.4 | -4.4 | -1.9 | -1.4 | -2.6 | -5.0 | -2.6 | -3.8 | 1.2 | 1.1 | 1.2 | 0.9 | 0.8 | 0.9 |
| 44 | FBgn0030556 | CG1810 | mRNA-cap | -2.4 | -2.2 | -1.8 | -4.4 | -1.4 | -1.9 | -2.6 | -4.4 | -2.0 | -3.3 | 0.9 | 0.9 | 1.1 | 0.7 | 0.5 | 0.7 |
| 45 | FBgn0029113 | CG7528 | Uba2 | -2.7 | -2.2 | 0.2 | -0.9 | -2.9 | -0.9 | -1.8 | -0.5 | 0.1 | -1.0 | 1.1 | 1.2 | 1.3 | 0.9 | 0.7 | 1.2 |
| 46 | FBgn0038585 | CG7993 | Non3 | -2.1 | -2.1 | 0.3 | -1.5 | -2.4 | -1.4 | -1.9 | 0.7 | 0.7 | 0.2 | 1.1 | 1.3 | 1.2 | 1.1 | 1.1 | 1.0 |
| 47 | FBgn0003612 | CG8068 | Su(var)2-10 | -3.4 | -3.9 | -3.1 | -1.3 | -1.3 | -1.9 | -2.2 | -0.8 | -0.4 | -1.5 | 1.5 | 1.3 | 1.3 | 1.3 | 1.3 | 1.0 |
| 48 | FBgn0032728 | CG18398 | Tango6 | -3.8 | -2.4 | -4.0 | -1.5 | -2.6 | -0.1 | -2.6 | -4.5 | -2.2 | -6.9 | 1.2 | 1.1 | 1.4 | 1.0 | 1.0 | 1.1 |
| 49 | FBgn0039406 | CG6090 | RpL34a | -4.6 | -8.7 | -1.1 | -4.5 | -2.4 | -1.8 | -0.6 | -6.9 | -5.6 | -8.1 | 2.4 | 2.3 | 2.1 | 1.5 | 1.4 | 1.9 |
| 50 | FBgn0263392 | CG2083 | Tet | -3.3 | -2.2 | 0.4 | 1.1 | -2.0 | -1.8 | -0.8 | -0.3 | 0.2 | -0.2 | 1.2 | 1.1 | 1.0 | 0.8 | 0.7 | 1.1 |
| 51 | FBgn0011205 | CG5725 | fbl | -2.4 | -2.1 | 0.3 | -1.6 | -1.6 | -1.6 | -0.5 | 1.3 | 0.1 | 1.3 | 1.0 | 1.0 | 1.0 | 0.8 | 0.7 | 1.1 |
| 52 | FBgn0027868 | CG6743 | Nup170 | -2.6 | -3.9 | -0.6 | -3.4 | -0.7 | -1.5 | -3.4 | -5.0 | -2.9 | -3.4 | 1.2 | 1.2 | 1.2 | 0.9 | 0.7 | 0.8 |
| General phagocytic factors | | | | | | | | | | | | | | | | | | | |
| 53 | FBgn0011742 | CG9901 | Arp14D | -11.6 | -3.0 | 0.9 | 0.0 | -3.1 | -2.1 | -4.6 | 0.6 | 1.3 | -0.1 | 0.5 | 0.4 | 0.5 | 0.3 | 0.3 | 0.3 |
| 54 | FBgn0025781 | CG6759 | cdc16 | -9.3 | -2.1 | -0.7 | -3.1 | -2.5 | -2.9 | -3.9 | -1.3 | -1.4 | -2.9 | 0.7 | 0.6 | 0.5 | 0.3 | 0.2 | 0.3 |
| 55 | FBgn0001205 | CG10367 | Hmgcr | -4.6 | -2.8 | -0.8 | -2.4 | -4.4 | -2.6 | -1.9 | -5.9 | -3.0 | -2.8 | 0.2 | 0.2 | 0.1 | 0.1 | 0.1 | 0.1 |
| 56 | FBgn0031437 | CG9881 | p16-ARC | -9.8 | -2.5 | 0.8 | -0.2 | -2.9 | -3.5 | -2.4 | 0.1 | 1.4 | 1.6 | 0.7 | 0.6 | 0.4 | 0.2 | 0.2 | 0.3 |
| 57 | FBgn0003660 | CG12210 | Syb | -3.9 | -2.5 | -1.6 | -0.1 | -1.6 | -1.5 | -3.6 | -2.1 | -0.3 | -1.1 | 0.4 | 0.3 | 0.4 | 0.2 | 0.3 | 0.4 |
| 58 | FBgn0035713 | CG10107 | velo | -3.4 | -4.8 | -0.5 | -5.5 | 0.3 | -2.4 | -3.4 | -1.7 | -0.7 | -1.7 | 0.7 | 0.6 | 0.7 | 0.7 | 0.7 | 0.5 |
| No human homologs | | | | | | | | | | | | | | | | | | | |
| 59 | FBgn0037127 | CG14566 | CG14566 | -2.8 | -2.3 | -0.7 | -0.4 | -4.2 | -3.8 | -4.1 | -4.0 | -0.2 | -1.5 | 1.6 | 1.9 | 1.7 | 1.3 | 1.2 | 1.5 |
| 60 | FBgn0033942 | CG10112 | Cpr51A | -5.8 | -2.1 | -2.2 | 0.4 | -0.7 | -1.6 | -1.9 | -0.6 | -0.1 | -0.2 | 1.0 | 1.0 | 1.0 | 0.9 | 0.7 | 0.9 |
| 61 | FBgn0025335 | CG4585 | CG4585 | -2.7 | -2.1 | -3.2 | -1.1 | 1.0 | -1.9 | -2.0 | -1.1 | 0.4 | 0.1 | 1.0 | 1.3 | 1.1 | 1.5 | 1.2 | 0.8 |
| Non-validated hits | | | | | | | | | | | | | | | | | | | |
| 62 | FBgn0027092 | CG10687 | Aats-asn | -4.2 | -2.7 | -0.3 | -1.1 | -1.2 | -1.3 | -2.7 | 0.7 | 1.9 | 0.3 | | | | | | |
| 63 | FBgn0033842 | CG5970 | cbc | -2.0 | -3.4 | -0.2 | -2.9 | -2.8 | -1.2 | -1.1 | -1.3 | -0.5 | -0.6 | | | | | | |
| 64 | FBgn0267861 | CG40196 | Maf1 | -3.7 | -2.2 | -1.8 | -1.2 | -0.8 | -1.1 | -2.8 | -0.5 | -0.1 | -1.2 | | | | | | |
| 65 | FBgn0050183 | CG30183 | CG30183 | -2.8 | -3.3 | 0.7 | -1.6 | -2.1 | -1.1 | -1.3 | -4.0 | -0.3 | -0.1 | | | | | | |
| 66 | FBgn0035726 | CG9953 | CG9953 | -2.2 | -2.2 | -2.2 | 0.8 | -1.0 | -1.9 | -1.3 | 0.8 | -0.2 | -0.4 | | | | | | |
| 67 | FBgn0010078 | CG3661 | RpL23 | -7.0 | -10.1 | -1.5 | -5.6 | -1.3 | -2.0 | -0.8 | -7.9 | -4.0 | -4.6 | | | | | | |
| 68 | FBgn0039923 | CG1793 | MED26 | -2.7 | -2.9 | -1.2 | -3.8 | -1.2 | -0.9 | -1.9 | -2.1 | -1.0 | -1.7 | | | | | | |
| 69 | FBgn0005674 | CG5394 | Aats-glupro | -3.9 | -4.4 | -0.8 | -3.4 | -3.1 | -0.5 | -0.4 | -2.6 | -0.2 | -0.2 | | | | | | |
| 70 | FBgn0022288 | CG12792 | l(2)09851 | -2.2 | -3.7 | 0.8 | -1.8 | -2.1 | -0.8 | -1.0 | 0.6 | 1.7 | 0.9 | | | | | | |
| 71 | FBgn0025800 | CG2262 | Smox | -2.4 | -2.7 | 0.1 | -1.4 | -0.7 | -1.7 | -0.8 | 0.6 | -0.6 | -0.6 | | | | | | |
| 72 | FBgn0051788 | CG31788 | CG31788 | -2.0 | -2.0 | -1.3 | -3.8 | -0.4 | -2.2 | -0.5 | -0.5 | -0.2 | 1.1 | | | | | | |
| 73 | FBgn0002524 | CG4162 | lace | -4.5 | -2.9 | -0.9 | 0.8 | -0.9 | -0.1 | -2.2 | 0.3 | 0.6 | -1.2 | | | | | | |
| 74 | FBgn0003525 | CG1395 | stg | -2.4 | -3.7 | -1.5 | -4.8 | -0.4 | -0.2 | -2.3 | -0.8 | 0.9 | -0.6 | | | | | | |
| 75 | FBgn0029799 | CG15772 | CG15772 | -6.7 | -3.5 | 0.6 | -2.7 | 0.2 | 0.2 | -3.0 | -0.8 | 0.3 | -0.5 | | | | | | |
| 76 | FBgn0050022 | CG30022 | CG30022 | -3.9 | -2.3 | -2.3 | -1.3 | -0.2 | -0.5 | -1.7 | -0.4 | -0.4 | -1.5 | | | | | | |
| 77 | FBgn0027783 | CG10212 | SMC2 | -4.2 | -2.9 | -1.9 | -2.7 | -2.1 | -0.3 | 0.1 | -0.3 | -0.7 | 0.5 | | | | | | |
| 78 | FBgn0027592 | CG4184 | MED15 | -3.5 | -2.6 | 0.7 | -1.2 | -1.6 | -0.4 | -0.2 | -1.7 | 0.5 | -0.1 | | | | | | |
| 79 | FBgn0025582 | CG9677 | Int6 | -8.8 | -6.1 | -1.6 | -4.2 | -0.3 | -2.2 | 0.6 | -6.9 | -4.1 | -6.6 | | | | | | |
| 80 | FBgn0002719 | CG10120 | Men | -2.5 | -2.0 | 1.2 | -1.0 | 0.6 | -1.6 | -0.7 | -0.2 | 0.7 | -0.7 | | | | | | |
| 81 | FBgn0036710 | CG6479 | CG6479 | -3.7 | -2.6 | 0.4 | -2.8 | 0.1 | -0.1 | -1.6 | -0.4 | 0.9 | -0.2 | | | | | | |
| 82 | FBgn0028691 | CG10230 | Rpn9 | -2.4 | -2.3 | -1.2 | -0.8 | 0.3 | -2.2 | 0.6 | 0.3 | -2.5 | -0.2 | | | | | | |
| 83 | FBgn0034461 | CG15124 | CG15124 | -2.5 | -2.4 | -0.6 | -2.9 | 1.0 | 0.1 | -1.8 | 0.1 | 0.5 | -1.4 | | | | | | |
| 84 | FBgn0036809 | CG12477 | CG12477 | -2.8 | -2.7 | -1.5 | -1.2 | -1.1 | -1.1 | -0.8 | -0.7 | 0.9 | 0.5 | | | | | | |
| 85 | FBgn0035120 | CG13879 | CG13879 | -2.2 | -2.3 | 0.0 | -1.9 | -1.4 | -1.2 | -0.4 | -0.6 | 0.0 | -0.1 | | | | | | |
| 86 | FBgn0029810 | CG12239 | CG12239 | -2.1 | -3.1 | 0.2 | -3.1 | -1.4 | -1.3 | 0.1 | 0.4 | 0.4 | -0.7 | | | | | | |
| 87 | FBgn0037294 | CG12590 | CG12590 | -2.9 | -2.1 | -0.8 | -3.2 | -1.4 | -0.2 | -0.8 | -0.5 | 0.0 | -0.1 | | | | | | |
| 88 | FBgn0037881 | CG10703 | CG10703 | -3.1 | -2.0 | -2.8 | -0.4 | -0.1 | -1.2 | -1.0 | -1.4 | 0.0 | -1.3 | | | | | | |
| 89 | FBgn0034933 | CG3735 | CG3735 | -2.1 | -2.1 | 1.4 | -0.6 | -0.4 | -1.4 | -0.3 | 0.4 | 1.3 | 1.5 | | | | | | |
| 90 | FBgn0001311 | CG2666 | kkv | -2.9 | -3.0 | 0.0 | -0.9 | -0.2 | -0.2 | -1.4 | -1.3 | 0.3 | -1.0 | | | | | | |
| 91 | FBgn0027617 | CG5808 | CG5808 | -2.0 | -3.1 | -1.5 | -4.6 | 0.2 | -0.7 | -1.1 | -0.7 | 0.5 | -1.2 | | | | | | |
| 92 | FBgn0037243 | CG9853 | CG9853 | -2.4 | -2.4 | -0.2 | 0.0 | -0.5 | -0.7 | -0.3 | 0.3 | 1.1 | 0.4 | | | | | | |
| 93 | FBgn0030306 | CG1751 | Spase25 | -2.1 | -2.7 | -1.4 | -2.8 | -0.7 | -0.3 | -0.2 | 0.7 | 1.2 | 1.1 | | | | | | |
| 94 | FBgn0036596 | CG13045 | CG13045 | -2.1 | -2.4 | -2.7 | -0.5 | -1.3 | -0.5 | 0.7 | 0.2 | 0.3 | 0.4 | | | | | | |
| 95 | FBgn0040290 | CG7487 | RecQ4 | -2.5 | -2.3 | -0.5 | -1.4 | 0.8 | -1.2 | -0.7 | 0.0 | 0.3 | -1.3 | | | | | | |
| 96 | FBgn0032404 | CG5317 | CG5317 | -2.5 | -3.9 | 0.3 | -0.4 | 0.1 | -0.3 | -0.8 | 0.8 | 0.6 | 0.3 | | | | | | |
| 97 | FBgn0042125 | CG18787 | CG18787 | -2.4 | -4.0 | 0.1 | -3.3 | 0.0 | -0.9 | 0.5 | 0.4 | 1.0 | 1.8 | | | | | | |
| 98 | FBgn0039966 | CG40304 | Rab21 | -2.1 | -3.7 | -1.1 | -2.3 | -0.1 | 0.0 | -0.3 | -0.2 | -0.1 | 0.3 | | | | | | |
| 99 | FBgn0038016 | CG10042 | MBD-R2 | -2.4 | -2.1 | -0.2 | -2.6 | 0.5 | -0.3 | -0.4 | -0.3 | 0.8 | 0.1 | | | | | | |
| 100 | FBgn0034056 | CG15699 | CG15699 | -4.4 | -3.1 | -0.5 | -3.5 | -1.2 | 1.1 | 0.0 | -0.5 | 0.0 | 0.6 | | | | | | |
| 101 | FBgn0032408 | CG6712 | CG6712 | -2.0 | -3.8 | 0.3 | -2.4 | -0.5 | 0.4 | 0.0 | 0.7 | 0.6 | 0.4 | | | | | | |
| 102 | FBgn0000617 | CG6474 | e(y)1 | -2.6 | -2.5 | -1.1 | -2.5 | 1.3 | -0.7 | -0.6 | 1.2 | -0.4 | 0.0 | | | | | | |
| 103 | FBgn0028398 | CG2859 | Taf10 | -2.5 | -2.2 | -1.7 | -0.3 | 0.4 | -0.2 | 0.0 | 0.0 | 0.8 | -0.1 | | | | | | |
| 104 | FBgn0033740 | CG8828 | CG8828 | -2.9 | -2.6 | -1.1 | -2.9 | 0.1 | 0.2 | 0.0 | -1.8 | 0.6 | 0.3 | | | | | | |
| 105 | FBgn0004860 | CG3895 | ph-d | -2.0 | -3.7 | -0.1 | -3.4 | -0.3 | 0.4 | 0.2 | 0.1 | 0.3 | -0.1 | | | | | | |
| 106 | FBgn0052000 | CG32000 | CG32000 | -3.4 | -2.9 | -0.7 | -2.4 | 0.5 | 0.8 | -0.6 | 0.0 | 1.1 | -1.7 | | | | | | |
| 107 | FBgn0052683 | CG32683 | CG32683 | -2.5 | -3.2 | -0.7 | -0.6 | 1.2 | -0.1 | -0.1 | 0.9 | 0.5 | -0.3 | | | | | | |
| 108 | FBgn0013953 | CG7005 | Esp | -4.2 | -2.7 | -3.9 | -0.9 | -0.1 | 0.7 | 0.5 | 0.8 | 1.2 | 1.1 | | | | | | |
| 109 | FBgn0026403 | CG12908 | Ndg | -2.6 | -2.3 | -4.1 | 0.7 | 0.4 | -0.1 | 0.9 | 0.4 | -2.9 | -0.2 | | | | | | |
| 110 | FBgn0035965 | CG14181 | Use1 | -2.7 | -2.5 | -0.7 | -1.6 | 0.9 | 1.5 | -0.8 | -1.9 | 0.4 | -2.4 | | | | | | |
| 111 | FBgn0038787 | CG4360 | CG4360 | -3.9 | -5.1 | -1.8 | -4.9 | 0.8 | 1.6 | 0.8 | -0.6 | 0.4 | -0.3 | | | | | | |
| 112 | FBgn0028835 | CG13383 | CSN8 | 2.5 | 2.1 | -0.7 | -3.5 | 0.7 | -2.5 | -0.3 | -4.7 | -2.6 | -1.5 | | | | | | |
| 113 | FBgn0005634 | CG15792 | zip | 2.4 | 2.4 | -1.7 | -3.5 | 0.1 | -0.6 | 0.5 | -4.4 | -2.0 | -3.2 | | | | | | |
| Low SL2 cell number in validation screen | | | | | | | | | | | | | | | | | | | |
| 114 | FBgn0003124 | CG12306 | polo | -8.9 | -3.1 | -1.5 | -3.5 | -4.4 | -4.9 | -4.1 | -9.4 | -6.4 | -7.1 | | | | | | |
| 115 | FBgn0032678 | CG12750 | ncm | -4.8 | -7.0 | -1.8 | -5.9 | -4.5 | -5.5 | -2.9 | -9.9 | -7.1 | -7.5 | | | | | | |
| 116 | FBgn0000405 | CG3510 | CycB | -2.9 | -3.3 | -2.3 | -1.0 | -2.1 | -3.9 | -3.1 | -11.4 | -6.5 | -10.1 | | | | | | |
| 117 | FBgn0025286 | CG1821 | RpL31 | -6.5 | -6.9 | -1.9 | -7.3 | -3.1 | -2.8 | -0.5 | -10.2 | -6.6 | -7.2 | | | | | | |
| 118 | FBgn0051155/FBgn0051344 | CG31155,CG31344 | CG31155, CG31344 | -3.5 | -2.3 | -1.7 | -6.1 | -1.4 | -1.7 | -3.1 | -9.1 | -7.5 | -9.3 | | | | | | |
| 119 | FBgn0002466 | CG10522 | sti | -2.4 | -2.9 | -0.9 | -3.6 | -1.3 | -0.5 | -0.9 | -9.3 | -6.1 | -7.0 | | | | | | |
| 120 | FBgn0003321 | CG1664 | sbr | -4.3 | -7.8 | -1.7 | -5.9 | -1.2 | -0.9 | -0.3 | -10.9 | -7.3 | -7.1 | | | | | | |
| Low SL2 cell number in first screen | | | | | | | | | | | | | | | | | | | |
| 121 | FBgn0035766 | CG8598 | eco | -5.7 | -2.9 | -3.5 | -2.0 | | | | | | | | | | | | |
| 122 | FBgn0035640 | CG17498 | mad2 | -4.1 | -3.8 | -1.9 | -4.3 | | | | | | | | | | | | |
| 123 | FBgn0016700 | CG3320 | Rab1 | -3.7 | -3.5 | -2.0 | -3.4 | | | | | | | | | | | | |
| 124 | FBgn0045862 | CG12878 | btz | -3.1 | -3.4 | -1.9 | -2.4 | | | | | | | | | | | | |
| 125 | FBgn0010280 | CG5444 | Taf4 | -2.6 | -5.6 | -2.0 | -5.2 | | | | | | | | | | | | |
| 126 | FBgn0030281 | CG11105 | CG11105 | -2.5 | -2.6 | -2.0 | -4.7 | | | | | | | | | | | | |
| 127 | FBgn0003862 | CG8651 | trx | -2.2 | -2.2 | -1.9 | -2.7 | | | | | | | | | | | | |
| 128 | FBgn0013746 | CG9556 | alien | 1.5 | 1.7 | -1.9 | -3.0 | | | | | | | | | | | | |
| 129 | FBgn0000100 | CG7490 | RpLP0 | -22.3 | -5.4 | -6.0 | -6.3 | | | | | | | | | | | | |
| 130 | FBgn0015288 | CG7434 | RpL22 | -16.0 | -5.1 | -5.7 | -5.4 | | | | | | | | | | | | |
| 131 | FBgn0034968 | CG3195 | RpL12 | -15.5 | -7.5 | -5.2 | -5.2 | | | | | | | | | | | | |
| 132 | FBgn0013325 | CG7726 | RpL11 | -14.5 | -6.0 | -6.2 | -4.5 | | | | | | | | | | | | |
| 133 | FBgn0036825 | CG6846 | RpL26 | -13.8 | -5.9 | -5.5 | -5.6 | | | | | | | | | | | | |
| 134 | FBgn0010409 | CG6510 | RpL18A | -13.4 | -6.3 | -5.1 | -5.5 | | | | | | | | | | | | |
| 135 | FBgn0035136 | CG6905 | CG6905 | -13.1 | -4.6 | -6.1 | -6.9 | | | | | | | | | | | | |
| 136 | FBgn0035753 | CG8615 | RpL18 | -12.2 | -5.9 | -5.1 | -4.9 | | | | | | | | | | | | |
| 137 | FBgn0029897 | CG3203 | RpL17 | -12.1 | -5.0 | -6.0 | -5.7 | | | | | | | | | | | | |
| 138 | FBgn0002607 | CG2746 | RpL19 | -11.9 | -5.1 | -4.8 | -5.1 | | | | | | | | | | | | |
| 139 | FBgn0000996 | CG8171 | dup | -11.7 | -4.6 | -4.7 | -7.0 | | | | | | | | | | | | |
| 140 | FBgn0005593 | CG4897 | RpL7 | -11.6 | -6.1 | -5.9 | -6.1 | | | | | | | | | | | | |
| 141 | FBgn0028688 | CG5378 | Rpn7 | -11.4 | -2.1 | -3.6 | -4.3 | | | | | | | | | | | | |
| 142 | FBgn0029785 | CG4111 | RpL35 | -11.1 | -5.5 | -4.3 | -4.9 | | | | | | | | | | | | |
| 143 | FBgn0002626 | CG7939 | RpL32 | -11.0 | -4.9 | -6.4 | -6.6 | | | | | | | | | | | | |
| 144 | FBgn0011272 | CG4651 | RpL13 | -10.9 | -5.9 | -2.3 | -3.4 | | | | | | | | | | | | |
| 145 | FBgn0038927 | CG6015 | CG6015 | -10.9 | -17.6 | -4.3 | -9.2 | | | | | | | | | | | | |
| 146 | FBgn0029856 | CG11700 | CG11700 | -10.5 | -3.3 | -5.7 | -7.3 | | | | | | | | | | | | |
| 147 | FBgn0037025 | CG11451 | CG11451 | -10.5 | -4.8 | -5.9 | -6.9 | | | | | | | | | | | | |
| 148 | FBgn0043044 | CG5887 | desat1 | -10.3 | -8.3 | -4.0 | -5.0 | | | | | | | | | | | | |
| 149 | FBgn0028968 | CG1528 | gammaCop | -10.1 | -3.7 | -6.0 | -5.7 | | | | | | | | | | | | |
| 150 | FBgn0030365 | CG1796 | Tango4 | -9.8 | -5.2 | -4.3 | -7.6 | | | | | | | | | | | | |
| 151 | FBgn0026872, FBgn0052808 | CG14777,CG32808 | CG14777, CG32808 | -9.5 | -9.2 | -2.6 | -4.5 | | | | | | | | | | | | |
| 152 | FBgn0031681, FBgn0261608 | CG31651,CG5827 | pgant5, RpL37A | -9.5 | -4.6 | -4.6 | -5.7 | | | | | | | | | | | | |
| 153 | FBgn0016697 | CG10938 | ProsMA5 | -9.5 | -4.7 | -5.3 | -6.9 | | | | | | | | | | | | |
| 154 | FBgn0040007 | CG40278 | RpL38 | -9.3 | -2.1 | -6.2 | -6.9 | | | | | | | | | | | | |
| 155 | FBgn0028692 | CG11888 | Rpn2 | -9.3 | -2.6 | -4.6 | -5.8 | | | | | | | | | | | | |
| 156 | FBgn0038834 | CG15697 | RpS30 | -9.3 | -4.5 | -4.8 | -6.6 | | | | | | | | | | | | |
| 157 | FBgn0003886 | CG9476 | alphaTub85E | -9.1 | -4.2 | -3.2 | -4.7 | | | | | | | | | | | | |
| 158 | FBgn0029134 | CG12323 | Prosbeta5 | -9.0 | -2.6 | -3.3 | -4.9 | | | | | | | | | | | | |
| 159 | FBgn0013749 | CG11027 | Arf102F | -8.9 | -5.7 | -6.3 | -6.4 | | | | | | | | | | | | |
| 160 | FBgn0015393 | CG3949 | hoip | -8.7 | -6.8 | -8.8 | -8.9 | | | | | | | | | | | | |
| 161 | FBgn0001218 | CG4147 | Hsc70-3 | -8.4 | -5.6 | -3.3 | -6.8 | | | | | | | | | | | | |
| 162 | FBgn0031980 | CG7424 | RpL36A | -8.4 | -4.4 | -2.9 | -5.6 | | | | | | | | | | | | |
| 163 | FBgn0026170 | CG4494 | smt3 | -8.2 | -2.5 | -5.9 | -6.4 | | | | | | | | | | | | |
| 164 | FBgn0030616 | CG9091 | RpL37a | -8.2 | -6.5 | -6.6 | -5.6 | | | | | | | | | | | | |
| 165 | FBgn0015756 | CG6141 | RpL9 | -7.9 | -6.3 | -6.0 | -3.9 | | | | | | | | | | | | |
| 166 | FBgn0001942 | CG9075 | eIF-4a | -7.9 | -7.4 | -5.8 | -4.8 | | | | | | | | | | | | |
| 167 | FBgn0004413 | CG10305 | RpS26 | -7.7 | -6.2 | -4.7 | -4.9 | | | | | | | | | | | | |
| 168 | FBgn0027936 | CG10682 | vih | -7.6 | -11.6 | -6.5 | -6.5 | | | | | | | | | | | | |
| 169 | FBgn0003517 | CG14792 | sta | -7.3 | -5.1 | -5.6 | -5.9 | | | | | | | | | | | | |
| 170 | FBgn0004378 | CG9191 | Klp61F | -7.2 | -6.9 | -4.4 | -6.1 | | | | | | | | | | | | |
| 171 | FBgn0003884 | CG1913 | alphaTub84B | -7.2 | -3.0 | -6.5 | -4.0 | | | | | | | | | | | | |
| 172 | FBgn0034084 | CG8435 | CG8435 | -7.2 | -6.8 | -4.5 | -4.4 | | | | | | | | | | | | |
| 173 | FBgn0010408 | CG3395 | RpS9 | -7.1 | -2.0 | -4.4 | -4.5 | | | | | | | | | | | | |
| 174 | FBgn0051611 | CG31611 | His4:CG31611 | -7.1 | -3.4 | -5.1 | -8.8 | | | | | | | | | | | | |
| 175 | FBgn0010590 | CG8392 | l(2)05070 | -7.0 | -3.1 | -6.1 | -5.5 | | | | | | | | | | | | |
| 176 | FBgn0028697 | CG40199 | | -6.9 | -10.3 | -3.0 | -6.6 | | | | | | | | | | | | |
| 177 | FBgn0061200 | CG4453 | Nup153 | -6.9 | -3.6 | -4.6 | -7.7 | | | | | | | | | | | | |
| 178 | FBgn0017579 | CG6253 | RpL14 | -6.7 | -10.0 | -5.2 | -6.5 | | | | | | | | | | | | |
| 179 | FBgn0001086 | CG4274 | fzy | -6.7 | -5.6 | -2.5 | -5.8 | | | | | | | | | | | | |
| 180 | FBgn0004404 | CG1527 | RpS14b | -6.7 | -2.9 | -2.8 | -5.5 | | | | | | | | | | | | |
| 181 | FBgn0015282 | CG5289 | Pros26.4 | -6.7 | -10.1 | -5.8 | -5.7 | | | | | | | | | | | | |
| 182 | FBgn0015745 | CG10652 | RpL30 | -6.6 | -5.6 | -5.5 | -4.3 | | | | | | | | | | | | |
| 183 | FBgn0033859 | CG6197 | CG6197 | -6.6 | -4.6 | -7.4 | -7.4 | | | | | | | | | | | | |
| 184 | FBgn0014189 | CG7269 | Hel25E | -6.6 | -9.0 | -5.8 | -3.9 | | | | | | | | | | | | |
| 185 | FBgn0031549 | CG15415 | CG15415 | -6.6 | -6.2 | -4.8 | -7.0 | | | | | | | | | | | | |
| 186 | FBgn0032518 | CG9282 | RpL24 | -6.6 | -5.3 | -5.2 | -4.9 | | | | | | | | | | | | |
| 187 | FBgn0000377 | CG3193 | crn | -6.5 | -4.3 | -3.9 | -8.3 | | | | | | | | | | | | |
| 188 | FBgn0010198 | CG2033 | RpS15Aa | -6.5 | -3.4 | -2.8 | -4.9 | | | | | | | | | | | | |
| 189 | FBgn0026380 | CG11981 | Prosbeta3 | -6.5 | -3.3 | -7.3 | -4.3 | | | | | | | | | | | | |
| 190 | FBgn0026664 | CG1515 | l(1)G0155 | -6.3 | -8.8 | -3.0 | -3.0 | | | | | | | | | | | | |
| 191 | FBgn0024733 | CG17521 | Qm | -6.2 | -12.6 | -6.0 | -5.5 | | | | | | | | | | | | |
| 192 | FBgn0064225 | CG17489 | RpL5 | -6.2 | -9.0 | -2.3 | -6.1 | | | | | | | | | | | | |
| 193 | FBgn0051613 | CG31613 | His3:CG31613 | -6.1 | -8.6 | -5.5 | -10.1 | | | | | | | | | | | | |
| 194 | FBgn0031266 | CG2807 | CG2807 | -5.8 | -6.3 | -5.7 | -6.5 | | | | | | | | | | | | |
| 195 | FBgn0039828 | CG1542 | CG1542 | -5.8 | -5.2 | -3.6 | -3.3 | | | | | | | | | | | | |
| 196 | FBgn0034908 | CG5543 | CG5543 | -5.7 | -3.6 | -4.6 | -5.4 | | | | | | | | | | | | |
| 197 | FBgn0025724 | CG6699 | beta'Cop | -5.7 | -6.3 | -7.0 | -6.0 | | | | | | | | | | | | |
| 198 | FBgn0019936 | CG15693 | RpS20 | -5.7 | -5.0 | -2.2 | -8.1 | | | | | | | | | | | | |
| 199 | FBgn0023170 | CG3997 | RpL39 | -5.6 | -4.4 | -4.7 | -6.1 | | | | | | | | | | | | |
| 200 | FBgn0261606, FBgn0053002 | CG15442, CG33002 | RpL27A, mRpL27 | -5.6 | -4.7 | -5.1 | -5.4 | | | | | | | | | | | | |
| 201 | FBgn0036548 | CG5931 | CG5931 | -5.5 | -13.4 | -4.3 | -8.4 | | | | | | | | | | | | |
| 202 | FBgn0010411 | CG8900 | RpS18 | -5.5 | -3.3 | -6.4 | -5.3 | | | | | | | | | | | | |
| 203 | FBgn0003701 | CG5785 | thr | -5.5 | -7.2 | -5.5 | -7.6 | | | | | | | | | | | | |
| 204 | FBgn0032987 | CG12775 | RpL21 | -5.4 | -10.8 | -2.8 | -7.5 | | | | | | | | | | | | |
| 205 | FBgn0015234 | CG8522 | HLH106 | -5.3 | -2.9 | -4.0 | -2.9 | | | | | | | | | | | | |
| 206 | FBgn0010265 | CG13389 | RpS13 | -5.3 | -6.1 | -3.7 | -4.8 | | | | | | | | | | | | |
| 207 | FBgn0015818 | CG3780 | Spx | -5.3 | -5.2 | -4.0 | -7.4 | | | | | | | | | | | | |
| 208 | FBgn0033912 | CG8415 | RpS23 | -5.2 | -2.9 | -4.6 | -3.4 | | | | | | | | | | | | |
| 209 | FBgn0036216 | CG14129 | CG14129 | -5.2 | -5.1 | -6.6 | -6.3 | | | | | | | | | | | | |
| 210 | FBgn0028695 | CG7762 | Rpn1 | -5.2 | -2.6 | -5.7 | -3.0 | | | | | | | | | | | | |
| 211 | FBgn0000986 | CG2637 | Fs(2)Ket | -5.1 | -4.7 | -3.0 | -4.3 | | | | | | | | | | | | |
| 212 | FBgn0025725 | CG7961 | alphaCop | -5.1 | -4.2 | -5.9 | -6.0 | | | | | | | | | | | | |
| 213 | FBgn0004856 | CG8264 | Bx42 | -5.0 | -4.9 | -6.2 | -8.0 | | | | | | | | | | | | |
| 214 | FBgn0017545 | CG2168 | RpS3A | -5.0 | -8.2 | -4.5 | -6.2 | | | | | | | | | | | | |
| 215 | FBgn0019949 | CG5179 | Cdk9 | -4.9 | -2.5 | -5.4 | -3.5 | | | | | | | | | | | | |
| 216 | FBgn0038947 | CG7073 | sar1 | -4.9 | -4.2 | -3.1 | -5.7 | | | | | | | | | | | | |
| 217 | FBgn0003279 | CG5502 | RpL4 | -4.9 | -14.7 | -4.7 | -6.1 | | | | | | | | | | | | |
| 218 | FBgn0002284 | CG4097 | Pros26 | -4.8 | -7.8 | -4.2 | -6.1 | | | | | | | | | | | | |
| 219 | FBgn0039154, FBgn0039154 | CG6164, CG6173 | Npc2f, Kal1 | -4.8 | -7.2 | -5.7 | -7.2 | | | | | | | | | | | | |
| 220 | FBgn0003600 | CG6476 | Su(var)3-9 | -4.8 | -7.1 | -4.7 | -4.7 | | | | | | | | | | | | |
| 221 | FBgn0037249 | CG9805 | eIF3-S10 | -4.8 | -8.9 | -2.9 | -6.3 | | | | | | | | | | | | |
| 222 | FBgn0033898 | CG8241 | CG8241 | -4.8 | -7.5 | -2.9 | -4.9 | | | | | | | | | | | | |
| 223 | FBgn0041147 | CG10850 | ida | -4.7 | -2.8 | -2.2 | -3.8 | | | | | | | | | | | | |
| 224 | FBgn0040227 | CG10161 | eIF-3p66 | -4.7 | -4.4 | -2.9 | -4.6 | | | | | | | | | | | | |
| 225 | FBgn0004177 | CG7109 | mts | -4.7 | -6.4 | -6.0 | -8.0 | | | | | | | | | | | | |
| 226 | FBgn0024998 | CG2685 | CG2685 | -4.6 | -3.7 | -2.6 | -5.5 | | | | | | | | | | | | |
| 227 | FBgn0029629 | CG8636 | CG8636 | -4.6 | -3.5 | -3.0 | -3.8 | | | | | | | | | | | | |
| 228 | FBgn0023167 | CG8427 | SmD3 | -4.6 | -5.8 | -2.2 | -6.4 | | | | | | | | | | | | |
| 229 | FBgn0027084 | CG12141 | Aats-lys | -4.6 | -4.6 | -3.7 | -4.5 | | | | | | | | | | | | |
| 230 | FBgn0039713 | CG7808 | RpS8 | -4.6 | -4.3 | -5.0 | -4.6 | | | | | | | | | | | | |
| 231 | FBgn0021760 | CG32435 | chb | -4.5 | -2.4 | -6.1 | -4.4 | | | | | | | | | | | | |
| 232 | FBgn0030150 | CG15321 | CG15321 | -4.5 | -4.3 | -2.2 | -4.2 | | | | | | | | | | | | |
| 233 | FBgn0013303 | CG7641 | Nca | -4.5 | -3.1 | -3.7 | -4.4 | | | | | | | | | | | | |
| 234 | FBgn0051755 | CG31755 | CG31755 | -4.5 | -2.5 | -3.8 | -3.9 | | | | | | | | | | | | |
| 235 | FBgn0010410 | CG15442 | RpL27A | -4.4 | -2.9 | -5.4 | -6.7 | | | | | | | | | | | | |
| 236 | FBgn0002590 | CG8922 | RpS5a | -4.4 | -7.4 | -2.4 | -5.0 | | | | | | | | | | | | |
| 237 | FBgn0003151 | CG4904 | Pros35 | -4.3 | -4.1 | -5.4 | -3.9 | | | | | | | | | | | | |
| 238 | FBgn0035946 | CG5651 | CG5651 | -4.2 | -6.7 | -2.1 | -6.1 | | | | | | | | | | | | |
| 239 | FBgn0002622 | CG6779 | RpS3 | -4.2 | -6.3 | -3.8 | -5.1 | | | | | | | | | | | | |
| 240 | FBgn0003941 | CG2960 | RpL40 | -4.2 | -7.8 | -2.9 | -6.9 | | | | | | | | | | | | |
| 241 | FBgn0034751 | CG3751 | RpS24 | -4.2 | -3.5 | -4.5 | -5.0 | | | | | | | | | | | | |
| 242 | FBgn0031467 | CG2973 | CG2973 | -4.2 | -4.1 | -3.4 | -3.5 | | | | | | | | | | | | |
| 243 | FBgn0036213 | CG7283 | RpL10Ab | -4.1 | -4.7 | -5.5 | -6.0 | | | | | | | | | | | | |
| 244 | FBgn0052603 | CG32603 | CG32603 | -4.1 | -7.1 | -2.7 | -4.2 | | | | | | | | | | | | |
| 245 | FBgn0028686 | CG16916 | Rpt3 | -4.1 | -5.9 | -3.3 | -3.5 | | | | | | | | | | | | |
| 246 | FBgn0004922 | CG10944 | RpS6 | -4.1 | -7.2 | -4.7 | -6.1 | | | | | | | | | | | | |
| 247 | FBgn0039626 | CG1420 | CG1420 | -4.0 | -5.8 | -4.3 | -6.6 | | | | | | | | | | | | |
| 248 | FBgn0003277 | CG1554 | RpII215 | -4.0 | -4.7 | -4.3 | -7.2 | | | | | | | | | | | | |
| 249 | FBgn0039680 | CG1911 | CAP-D2 | -4.0 | -2.7 | -2.4 | -3.1 | | | | | | | | | | | | |
| 250 | FBgn0035648 | CG13288 | CG13288 | -3.9 | -2.7 | -3.2 | -5.9 | | | | | | | | | | | | |
| 251 | FBgn0010417 | CG32211 | Taf6 | -3.9 | -2.7 | -4.3 | -5.3 | | | | | | | | | | | | |
| 252 | FBgn0039757 | CG1883 | RpS7 | -3.9 | -9.6 | -2.9 | -5.7 | | | | | | | | | | | | |
| 253 | FBgn0015521 | CG2986 | oho23B | -3.8 | -3.8 | -3.5 | -4.8 | | | | | | | | | | | | |
| 254 | FBgn0015618 | CG10572 | Cdk8 | -3.8 | -4.3 | -5.2 | -5.1 | | | | | | | | | | | | |
| 255 | FBgn0033902 | CG8309 | Tango7 | -3.8 | -4.3 | -2.9 | -3.5 | | | | | | | | | | | | |
| 256 | FBgn0027049 | CG7769 | DDB1 | -3.8 | -5.2 | -2.1 | -3.4 | | | | | | | | | | | | |
| 257 | FBgn0014163 | CG4609 | fax | -3.8 | -3.3 | -4.2 | -2.2 | | | | | | | | | | | | |
| 258 | FBgn0011640 | CG8597 | lark | -3.8 | -5.5 | -2.7 | -3.5 | | | | | | | | | | | | |
| 259 | FBgn0004379 | CG10923 | Klp67A | -3.8 | -3.4 | -3.1 | -3.8 | | | | | | | | | | | | |
| 260 | FBgn0040394 | CG16903 | CG16903 | -3.7 | -6.7 | -2.1 | -4.2 | | | | | | | | | | | | |
| 261 | FBgn0033400 | CG2063 | CG2063 | -3.6 | -3.9 | -2.8 | -6.3 | | | | | | | | | | | | |
| 262 | FBgn0020910 | CG4863 | RpL3 | -3.6 | -7.3 | -4.9 | -5.6 | | | | | | | | | | | | |
| 263 | FBgn0038464 | CG16941 | CG16941 | -3.6 | -6.0 | -4.0 | -4.8 | | | | | | | | | | | | |
| 264 | FBgn0037686 | CG9354 | RpL34b | -3.6 | -4.0 | -3.8 | -4.9 | | | | | | | | | | | | |
| 265 | FBgn0014010 | CG3664 | Rab5 | -3.5 | -2.1 | -4.8 | -3.8 | | | | | | | | | | | | |
| 266 | FBgn0033688 | CG8877 | prp8 | -3.4 | -5.7 | -2.4 | -8.1 | | | | | | | | | | | | |
| 267 | FBgn0014127 | CG10726 | barr | -3.4 | -2.4 | -4.2 | -5.0 | | | | | | | | | | | | |
| 268 | FBgn0020255 | CG1404 | ran | -3.4 | -7.0 | -3.7 | -5.4 | | | | | | | | | | | | |
| 269 | FBgn0030343 | CG1886 | ATP7 | -3.4 | -4.9 | -3.1 | -4.0 | | | | | | | | | | | | |
| 270 | FBgn0005533 | CG3922 | RpS17 | -3.3 | -9.7 | -3.7 | -8.4 | | | | | | | | | | | | |
| 271 | FBgn0028694 | CG18174 | Rpn11 | -3.3 | -6.6 | -3.4 | -4.8 | | | | | | | | | | | | |
| 272 | FBgn0011217 | CG7425 | eff | -3.2 | -3.5 | -4.2 | -2.9 | | | | | | | | | | | | |
| 273 | FBgn0004867 | CG5920 | sop | -3.2 | -3.3 | -4.9 | -4.8 | | | | | | | | | | | | |
| 274 | FBgn0040477 | CG13329 | cid | -3.2 | -4.5 | -2.7 | -3.0 | | | | | | | | | | | | |
| 275 | FBgn0014366 | CG2925 | noi | -3.1 | -4.7 | -4.0 | -4.6 | | | | | | | | | | | | |
| 276 | FBgn0028411 | CG12752 | Nxt1 | -3.1 | -3.6 | -4.9 | -6.5 | | | | | | | | | | | | |
| 277 | FBgn0010355 | CG17603 | Taf1 | -3.0 | -2.6 | -2.0 | -3.6 | | | | | | | | | | | | |
| 278 | FBgn0031962 | CG18591 | CG18591 | -3.0 | -5.4 | -5.3 | -4.5 | | | | | | | | | | | | |
| 279 | FBgn0001491 | CG1639 | l(1)10Bb | -2.9 | -7.5 | -2.2 | -6.9 | | | | | | | | | | | | |
| 280 | FBgn0024188 | CG12352 | san | -2.9 | -3.7 | -2.0 | -4.0 | | | | | | | | | | | | |
| 281 | FBgn0035205 | CG2469 | CG2469 | -2.9 | -2.0 | -3.2 | -3.0 | | | | | | | | | | | | |
| 282 | FBgn0010083 | CG5352 | SmB | -2.9 | -6.6 | -5.1 | -6.3 | | | | | | | | | | | | |
| 283 | FBgn0028969 | CG14813 | deltaCOP | -2.9 | -7.6 | -3.7 | -7.5 | | | | | | | | | | | | |
| 284 | FBgn0038760 | CG12254 | MED25 | -2.9 | -2.1 | -4.6 | -4.0 | | | | | | | | | | | | |
| 285 | FBgn0033795 | CG17054 | Cap-G | -2.8 | -2.1 | -2.2 | -5.7 | | | | | | | | | | | | |
| 286 | FBgn0035210 | CG2213 | CG2213 | -2.8 | -2.9 | -2.4 | -2.6 | | | | | | | | | | | | |
| 287 | FBgn0027567 | CG8108 | CG8108 | -2.7 | -6.2 | -2.8 | -4.1 | | | | | | | | | | | | |
| 288 | FBgn0024921 | CG7398 | Trn | -2.7 | -3.1 | -2.6 | -2.5 | | | | | | | | | | | | |
| 289 | FBgn0035145 | CG12031 | MED14 | -2.7 | -3.1 | -5.2 | -4.9 | | | | | | | | | | | | |
| 290 | FBgn0013969 | CG5519 | Gbp | -2.7 | -3.4 | -2.8 | -4.7 | | | | | | | | | | | | |
| 291 | FBgn0011703 | CG5371 | RnrL | -2.6 | -1.9 | -6.3 | -7.7 | | | | | | | | | | | | |
| 292 | FBgn0032634 | CG6840 | Rpb11 | -2.6 | -4.5 | -3.6 | -6.2 | | | | | | | | | | | | |
| 293 | FBgn0038667 | CG5454 | CG5454 | -2.6 | -2.9 | -5.5 | -5.3 | | | | | | | | | | | | |
| 294 | FBgn0005648 | CG2163 | Pabp2 | -2.6 | -2.0 | -2.1 | -5.2 | | | | | | | | | | | | |
| 295 | FBgn0020277 | CG8807 | lush | -2.5 | -2.3 | -2.7 | -3.5 | | | | | | | | | | | | |
| 296 | FBgn0011771 | CG5837 | Hem | -2.5 | -2.2 | -3.7 | -2.2 | | | | | | | | | | | | |
| 297 | FBgn0031886 | CG8902 | CG8902 | -2.5 | -2.3 | -2.4 | -2.4 | | | | | | | | | | | | |
| 298 | FBgn0004855 | CG3284 | RpII15 | -2.4 | -3.2 | -5.2 | -5.4 | | | | | | | | | | | | |
| 299 | FBgn0031035 | CG14206 | RpS10b | -2.4 | -5.6 | -4.1 | -5.7 | | | | | | | | | | | | |
| 300 | FBgn0015521, FBgn0031473 | CG2986, CG3104 | RpS21, CG3104 | -2.3 | -5.1 | -4.1 | -5.3 | | | | | | | | | | | | |
| 301 | FBgn0037434 | CG1249 | CG1249 | -2.3 | -6.3 | -4.3 | -5.7 | | | | | | | | | | | | |
| 302 | FBgn0003276 | CG3180 | RpII140 | -2.2 | -3.9 | -4.3 | -5.1 | | | | | | | | | | | | |
| 303 | FBgn0003963 | CG2762 | ush | -2.2 | -7.8 | -4.9 | -7.7 | | | | | | | | | | | | |
| 304 | FBgn0033156 | CG12165 | Incenp | -2.2 | -2.4 | -6.1 | -6.9 | | | | | | | | | | | | |
| 305 | FBgn0037357 | CG1250 | sec23 | -2.2 | -6.2 | -2.5 | -6.7 | | | | | | | | | | | | |
| 306 | FBgn0027603 | CG12359 | Ulp1 | -2.1 | -3.0 | -2.7 | -4.4 | | | | | | | | | | | | |
| 307 | FBgn0032596 | CG17331 | CG17331 | -2.1 | -3.8 | -6.4 | -5.4 | | | | | | | | | | | | |
| 308 | FBgn0034926 | CG5591 | CG5591 | -2.1 | -3.8 | -4.1 | -3.1 | | | | | | | | | | | | |
| 309 | FBgn0003275 | CG1163 | RpII18 | -2.1 | -4.3 | -6.0 | -6.4 | | | | | | | | | | | | |
| 310 | FBgn0031771 | CG9140 | CG9140 | -2.1 | -2.9 | -2.8 | -2.1 | | | | | | | | | | | | |
| 311 | FBgn0027571 | CG3523 | CG3523 | -2.0 | -6.6 | -3.7 | -5.8 | | | | | | | | | | | | |
| 312 | FBgn0010621 | CG8439 | Cct5 | -2.0 | -4.3 | -3.5 | -3.6 | | | | | | | | | | | | |
| 313 | FBgn0039830 | CG1746 | CG1746 | -2.0 | -3.3 | -3.5 | -4.7 | | | | | | | | | | | | |
